# A congenital hydrocephalus causing mutation in Trim71 results in stem cell differentiation defects through inhibiting *Lsd1* mRNA translation

**DOI:** 10.1101/2022.04.14.488304

**Authors:** Qiuying Liu, Mariah K. Novak, Rachel M. Pepin, Katharine R. Maschhoff, Xiaoli Chen, Shaojie Zhang, Wenqian Hu

## Abstract

Congenital hydrocephalus (CH) is a major cause of childhood morbidity. Mono-allelic mutations in Trim71, a conserved stem-cell-specific RNA-binding protein, cause CH, however, molecular basis for pathogenesis mediated by these mutations remains unknown. Here, using mouse embryonic stem cells as a model, we reveal that the mouse R783H mutation (R796H in human) significantly alters Trim71’s mRNA substrate specificity and leads to accelerated stem-cell differentiation and neural lineage commitment. The mutant Trim71, but not the wild-type Trim71, binds *Lsd1 (Kdm1a)* mRNA and represses its translation. Specific inhibition of this repression or a slight increase of Lsd1 in the mutant cells alleviates the defects in stem cell differentiation and neural lineage commitment. These results determine a functionally relevant target of the CH-causing Trim71 mutant that can potentially be a therapeutic target and provide molecular mechanistic insights into the pathogenesis of this disease.

## Introduction

RNA-binding proteins (RBPs) control mRNA fate and are critical regulators of gene expression (Glisovic *et al*, 2008). These proteins play essential roles in animal development, and aberrations in RBPs contribute to a wide variety of human diseases (Brinegar & Cooper, 2016; Gebauer *et al*, 2021; Lukong *et al*, 2008). While the details of RBP-mediated regulations in a diverse range of physiological processes are becoming increasingly clear, our understanding of the molecular mechanisms by which mutations in RBPs result in diseases is still limited.

Congenital hydrocephalus (CH), a significant cause of childhood morbidity, is caused by imbalanced neurogenesis that leads to abnormal accumulation of cerebrospinal fluid in brain ventricles (Kahle *et al*, 2016). The primary treatment for CH is neurosurgical shunting, which has numerous complications. Human genome-wide association studies and pedigree analysis have identified several CH-causing mutations, including two mono-allelic and potential gain-of-function missense mutations in Trim71 (Furey *et al*, 2018; Jin *et al*, 2020), a stem-cell specific RBP that is highly conserved from *C. elegans* to human (Connacher & Goldstrohm, 2021; Ecsedi & Grosshans, 2013).

Trim71 binds to target mRNAs and down-regulates their expression through translational repression and/or enhanced mRNA degradation (Aeschimann *et al*, 2017; Chang *et al*, 2012; Liu *et al*, 2021a; Loedige *et al*, 2013; Welte *et al*, 2019; Worringer *et al*, 2014). Genetic studies in mice indicate that Trim71 is essential for early embryogenesis and proper neural differentiation, indicating it has critical functions during normal development (Chen *et al*, 2012; Cuevas *et al*, 2015; Maller Schulman *et al*, 2008). Mice with the homologous human CH-causing point mutations in Trim71 also have CH and display defects in neurogenesis (Duy *et al*, 2022), arguing for conserved mechanisms of pathogenesis from the Trim71 mutants. How these mutations in Trim71 cause diseases, such as CH, however, is still unknown.

Here, we dissected the molecular mechanisms of pathogenesis mediated by a disease-causing mutation in Trim71. Using mouse embryonic stem cells (mESCs) as a model, we compared the transcriptome-wide targets of wide-type (WT) Trim71 and the Trim71 bearing a homologous human CH-causing mutation (R783H in mouse, which is equivalent to R796H in human). This mutation significantly alters Trim71’s mRNA substrates and leads to accelerated stem cell differentiation and neural lineage commitment. Among the mRNAs that are uniquely bound by the mutant Trim71, we determined that the mutant Trim71 represses *Lsd1* (*Kdm1a*) mRNA translation. Specific inhibition of this repression through deleting the mutant Trim71’s binding site in the 3’UTR of *Lsd1* mRNA or a modest increase of Lsd1 in the mutant mESCs alleviates the defects in stem cell differentiation and neural lineage commitment. Altogether, these results determine a functionally relevant target of the CH-causing Trim71 mutant that can potentially be a therapeutic target and provide molecular mechanistic insights into the pathogenesis of CH caused by the R796H mutation in *TRIM71*. Moreover, our finding revealed that a disease-causing mutation in an RBP does not abolish RNA binding but alters its binding specificity, a mechanism by which gain-of-function mutations in RBPs can result in disease.

## Results

### R783H Trim71 causes stem-cell and neural-differentiation defects

We used mESCs as a model to study the CH-causing mutations in Trim71 because: a) Trim71 is a highly conserved stem-cell-specific RBP (Connacher & Goldstrohm, 2021; Ecsedi & Grosshans, 2013); b) the homologous human CH-causing point mutations in mouse Trim71 also lead to CH and neurogenesis defects (Duy *et al*., 2022). We chose the FLAG-Trim71 mESCs for mechanistic studies, because the bi-allelic FLAG-tag knock-in at the N-terminus of Trim71 in these mESCs enables unambiguous detection and isolation of the endogenous Trim71 using an anti-FLAG monoclonal antibody (Liu *et al*., 2021a). Here, we studied the R783H Trim71 mutation in mouse, which is equivalent to the CH-causing R796H mutation in human TRIM71 (Furey *et al*., 2018; Jin *et al*., 2020). This mutation is within the RNA-binding domain of Trim71 (Figure 1-figure supplement 1A), suggesting an alteration of the interactions between Trim71 and its target RNAs.

**Figure 1.**
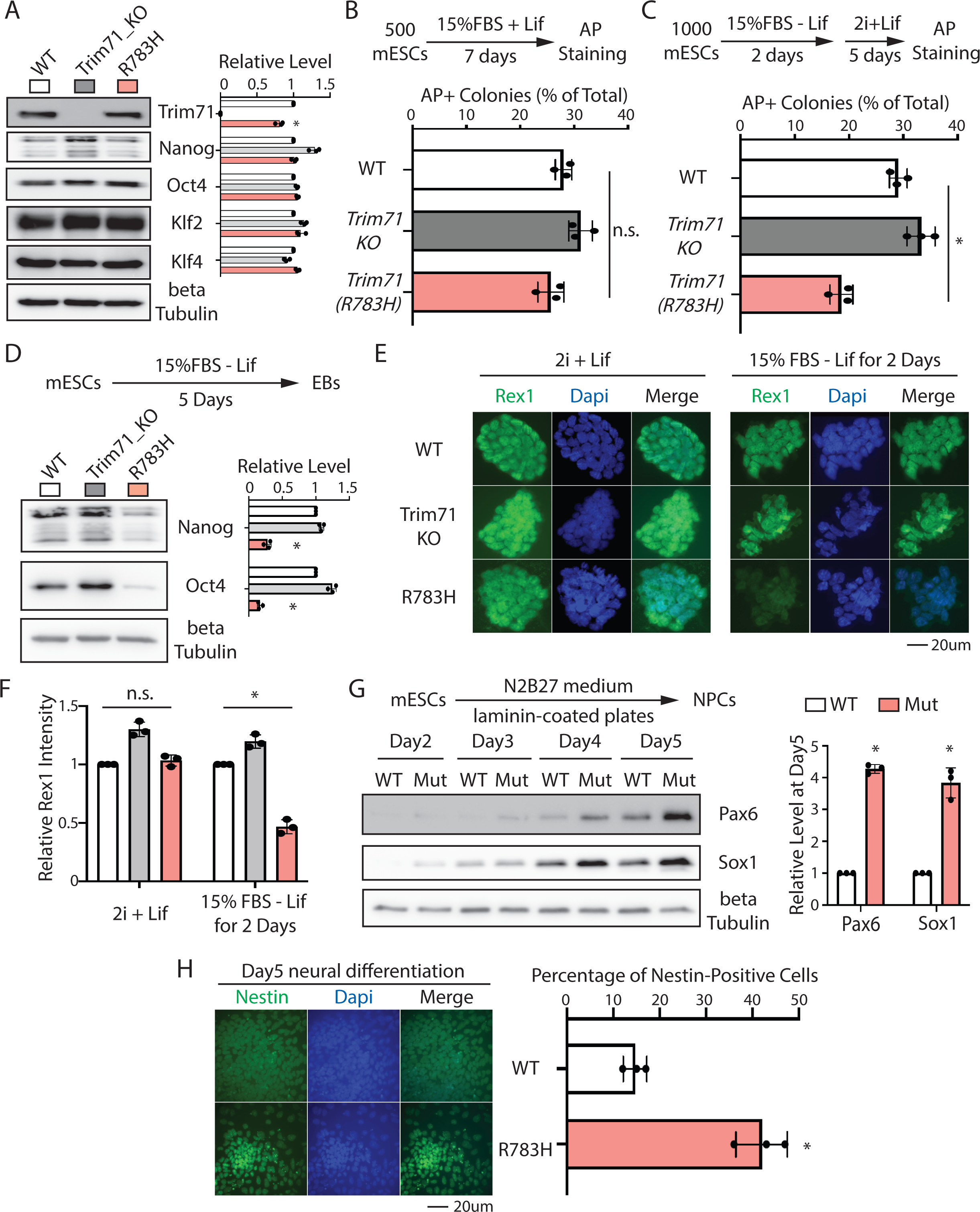
The CH-causing Trim71 mutation results in stem cell and neural differentiation defects in mESCs. A. Western blotting in the WT, Trim71 knockout (Trim71_KO), and the Trim71 mutant (R783H) mESCs. B. Colony formation assay for mESCs. The mESCs were cultured in 15%FBS + Lif for 7 days, and the resultant colonies were fixed and stained for AP. C. Exit pluripotency assay for mESCs. The mESCs were induced to exit pluripotency in medium without Lif for 2 days and then switched to 2i+Lif medium for 5 days. The resultant colonies were fixed and stained for AP. D. Western blotting of pluripotency factors during EB formation. E. Immunofluorescence (IF) staining showing the expression level of Rex1 in mESCs cultured in the stemness (2i + Lif) and differentiating (15% FBS – Lif) conditions. F. Relative intensity of IF signals from individual cells in experiment associated with E. G. Expression of neural lineage markers during mESCs neural differentiation. H. IF staining of Nestin. The quantifications represent the means (± SD) of three independent experiments. In A, D, and G, representative Western blots are shown, and the quantifications represent the means (± SD) of three independent experiments. In B and C, the colony morphology and AP intensity were evaluated through microscopy. 100-200 colonies were examined each time to determine the percentage of undifferentiated colonies. The results represent the means (± SD) of three independent experiments. Student’s t-test was used to determine the significance of the difference, *p<0.05; and n.s. not significant (p>0.05). The following figure supplements are available for Figure 1: Figure supplement 1. Generation of the Trim71 R783H monoallelic and bi-allelic mESCs. Figure supplement 2. The R783H mutation in Trim71 does not change the proliferation and apoptosis of mESCs. Figure supplement 3. The R783H mutation in Trim71 does not alter the microRNA pathway in mESCs. Figure supplement 4. The R783H mESCs are more prone differentiate into the ectoderm lineage during EB formation. Figure 1 – source data Tiff files of raw gel images for Figure 1A, D, G; Figure 1-figure supplement 3.

Using genome editing, we generated both monoallelic and bi-allelic R783H mutations on Trim71 in the FLAG-Trim71 mESCs (Figure 1-figure supplement 1B). In human, the monoallelic R796H mutation in TRIM71 causes CH, arguing that this mutation can be a gain-of-function mutation (Furey *et al*., 2018; Jin *et al*., 2020). A challenge of mechanistic studies in the heterozygous background, however, is that it is difficult to discriminate whether the identified phenotypes/interactions (e.g., mRNA substrates) are mediated directly by the mutant protein or by the potential alterations of the WT protein (e.g., potential dimerization between WT and mutant proteins). To circumvent this, we first used mESCs with the bi-allelic mutation for functional and mechanistic studies on the R783H Trim71 mutant, and then we examined whether the identified mechanistic insights are disease relevant in mESCs with the monoallelic R783H mutation in Trim71.

The R783H mutation, in either the homozygous or the heterozygous background, does not alter the proliferation and apoptosis of mESCs under both stemness and differentiating conditions (Figure 1-figure supplement 2). In contrast, the Trim71 knockout (KO) mESCs displayed impaired growth and increased apoptosis (Figure 1- figure supplement 2), which is consistent with the previous report (Chang *et al*., 2012). These results argue that the R783H is not a loss-of-function mutation. Moreover, the R783H mutation does not impact the microRNA pathway in mESCs, because neither the levels of Ago2, the major argonaute protein in mESCs (Liu *et al*, 2021b), nor a group of microRNAs involved in either differentiation (e.g., let-7 microRNAs) or pluripotency (e.g., the miR-290, 291, 293) are altered in the R783H mutant mESCs (Figure 1-figure supplement 3).

In mESCs with the bi-allelic mutation, the R783H mutation resulted in a modest decrease (∼80% of WT levels) in Trim71 levels (Figure 1A) but did not impact stem cell self-renewal, as revealed by either examining the expression of pluripotency factors (Figure 1A) or colony formation assays (Figure 1B). However, when subjected to the exit pluripotency assay, which evaluates the rate mESCs exit the pluripotent state (Betschinger *et al*, 2013), the R783H mutant mESCs lost pluripotency at a significantly increased rate compared to either the WT or the Trim71 knockout mESCs (Figure 1C). Moreover, when subjected to differentiation via embryonic body (EB) formation for 5 days, the R783H mutant mESCs showed decreased levels of pluripotency factors than either the WT or Trim71 knockout mESCs (Figure 1D). Consistent with these findings, immunofluorescence staining revealed that differentiating R783H mutant mESCs had less Rex1, a marker of pluripotency, than either the WT or the Trim71 KO mESCs (Figure 1E&F). These results collectively indicate that the R783H mutant mESCs are more prone to differentiation and argue that the R783H Trim71 mutation is a gain-of-function mutation, which is consistent with the observations that monoallelic R796H mutation in TRIM71 results in CH (Furey *et al*., 2018).

As CH is a neurological disorder (Kahle *et al*., 2016), we subjected mESCs to monolayer neural differentiation (Mulas *et al*, 2019) and monitored the appearance of neural progenitor cells by examining the expression of Sox1 and Pax6, two critical transcription factors essential for neuroectodermal specification in mammals (Li *et al*, 2005). Compared to the WT mESCs, R783H mESCs showed both the earlier appearance and increased immunoblotting intensity of these two factors during the neural differentiation (Figure 1G), indicating that the CH-causing R783H Trim71 mutation resulted in accelerated neural differentiation in mESCs. Consistently, immunofluorescence staining of Nestin, a marker of neural progenitor cells, indicated the R783H cells express higher Nestin than the WT cells at the Day5 of neural differentiation (Figure 1H). Moreover, when subjected to spontaneous differentiation through EB formation, the R783H mESCs specifically expressed more ectoderm markers than the WT mESCs (Figure 1-figure supplement 4). Altogether, these results indicated that the R783H Trim71 mutant led to stem cell and neural differentiation defects in mESCs.

### R783H Trim71 has altered target mRNA binding

The R783H mutation is located in the RNA-binding domain of Trim71 (Figure 1-figure supplement 1A). To determine how this mutation impacts Trim71:RNA interactions, we identified transcriptome-wide targets of the R783H Trim71 mutant in mESCs using crosslinking immunoprecipitation and sequencing (CLIP-seq) (Figure 2A) (Darnell, 2010), which also revealed the R783H Trim71 binding region(s) on the target mRNAs. CLIP-seq was performed on the mESCs grown in the 2i+lif medium, which suppresses differentiation and maintains mESCs in the ground state (Ying *et al*, 2008). These culture conditions eliminated the potential differentiation differences between the WT and the R783H mESCs (Figure 2-figure supplement 1) and enabled us to evaluate how the CH-causing mutation impacts Trim71’s target recognition under the same developmental state.

**Figure 2.**
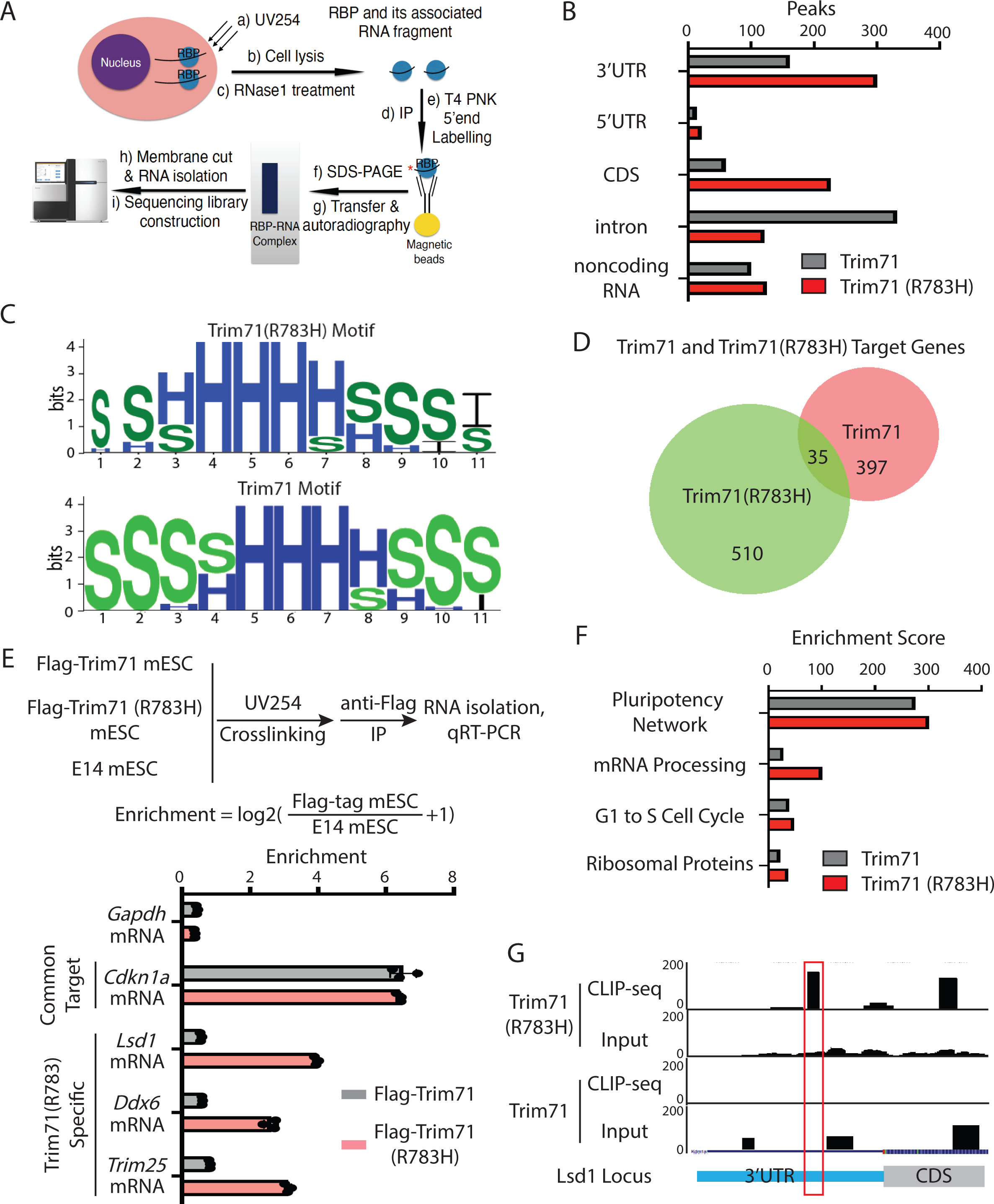
Transcriptome-wide identification of the target mRNAs of the Trim71 mutant R783H in mESCs. A. A work flow of the CLIP-seq analysis. B. Distribution of the WT Trim71 and the R783H Trim71 mutant binding regions in the mouse genome. In each CLIP-seq, there are two biological replicates. The binding regions present in both of the biological replicates are used for the analysis. C. Comparison of RNA secondary structures over-represented in the Trim71 mutant R783H and Trim71. “H”, “S”, and “I” indicates a nucleotide in a hairpin loop region, a stack region, and an internal loop region, respectively. D. Venn diagram showing the genes with binding sites from the WT Trim71 and the Trim71 mutant R783H. E. CLIP-qRT-PCR for the identified target mRNAs of the Trim71 mutant R783H. The results represent the means (±SD) of three independent experiments. F. Gene ontology analysis of the mRNAs with 3’UTR binding sites from the WT Trim71 and the R783H Trim71 mutant. G. UCSC genome browser snapshot for the CLIP-seq data from Trim71 and the Trim71 mutant R783H in the *Lsd1* locus. The red box indicates the binding region of the Trim71 mutant R783H. The inputs are from the size-matched input samples in the CLIP-seq analysis. The following figure supplement is available for Figure 2: Figure supplement 1. Colony formation assay for the WT and the R783H mESCs grown in the 2i+lif medium. Figure supplement 2. RNA-seq analysis of the WT and R783H mESCs. Figure supplement 3. Identification of potential functional targets of the R783H Trim71 mutant. Figure 2 - source data Tiff files of raw gel images for Figure 2-figure supplement 3.

Comparative analysis of the CLIP-seq data from R783H Trim71 and WT Trim71 (Liu *et al*., 2021a) revealed two similarities. First, 3’UTR is one of the major binding regions for both the WT and the mutant Trim71 (Figure 2B, Supplementary file1).

Second, WT and mutant-binding sites have a similar over-representation of predicted stem-loop structures, but no enriched primary sequence motifs, compared to randomized sequences, consistent with the results from *in vitro* studies that Trim71 recognizes structural motifs (Kumari *et al*, 2018) (Figure 2C). Despite these common features, there is only a small overlap between the mRNAs bound by WT Trim71 and the mutant Trim71 (Figure 2D), implying that the mutant Trim71 regulates a different set of mRNAs compared to WT Trim71 does. To validate this finding, we performed CLIP-qRT-PCR in WT and the mutant mESCs. Both WT and the mutant Trim71 bound *Cdkn1a* mRNA, a common target identified in the CLIP-seq data, however, only the mutant Trim71 bound *Lsd1*, *Ddx6*, and *Trim25* mRNAs (Figure 2E), three of the mutant-specific targets identified in the CLIP-seq data. Altogether, these results indicated that the CH-causing mutation in Trim71 significantly alters the substrate mRNAs to which it binds.

To determine whether or not this alteration of substrate RNAs is due to difference of RNA availability, we surveyed the transcriptomes of the WT and the R783H mutant mESCs grown in the 2i+lif medium, where both of these two types of mESCs have the same developmental status (Figure 2-figure supplement 1). Most (541 out of 545) of the R783H Trim71 mutant’s target RNAs were not differentially expressed between the WT and the R783H mESCs (Figure 2-figure supplement 2), indicating that the difference of substrate RNAs between the WT and the R783H mutant Trim71 is not caused by the RNA availability in the WT and the R783H mESCs. Moreover, this result also argues that the R783H Trim71 mutant does not destabilize its substrate RNAs.

Gene ontology analysis revealed that mRNAs specifically bound by the R783H Trim71 mutant were over-represented for the genes involved in regulating stem cell differentiation and pluripotency (Figure 2F), consistent with the finding that the R783H Trim71 mutant mESCs displayed stem cell and neural differentiation defects (Figure 1). A caveat in interpreting these results is that binding does not necessarily result in expression changes. To identify the functional targets of the R783H Trim71 mutant, we used the following criteria: a) mRNAs uniquely bound by the mutant Trim71 but not the WT Trim71; b) mRNAs encoding proteins with conserved functions in controlling stem cell differentiation; c) mRNAs abundantly expressed in mESCs. Western blotting on the resulting candidates revealed that Lsd1 consistently had the most decreased levels in the mutant mESCs compared to WT mESCs (Figure 2-figure supplement 3), arguing that *Lsd1* mRNA may be a functional target of the R783H Trim71 mutant. Lsd1 (Kdma1) is a conserved lysine-specific histone demethylase that is critical for both pluripotency and neural lineage commitment (Han *et al*, 2014; Whyte *et al*, 2012). The CLIP-seq data indicated that there is a R783H mutant Trim71 specific binding peak in the 3’UTR of *Lsd1* mRNA, and this peak’s signal is significantly higher than that from the size-matched input control (Van Nostrand *et al*, 2016) (Figure 2G), indicating that in mESCs, the mutant Trim71, but not the WT Trim71, specifically interacts with this region of *Lsd1* mRNA. In the subsequent experiments, we focused on the interaction between *Lsd1 (Kdm1a)* mRNA and the mutant Trim71.

### R783H Trim71 represses *Lsd1* mRNA translation

Multiple lines of evidence indicated that the R783H Trim71 mutant represses *Lsd1* mRNA translation in mESCs. First, Lsd1 protein decreased ∼2 fold with no significant changes in the level of *Lsd1* mRNA in mutant mESCs compared to WT mESCs (Figure 3A&B). Second, polysome analysis, which examines mRNA and ribosome association, revealed that *Lsd1* mRNA, but not a control mRNA, is translationally repressed in mutant mESCs compared to WT mESCs (Figure 3C&D). Third, when ectopically expressed in the WT mESCs, the R783H Trim71 mutant, but not WT Trim71, decreased Lsd1 protein levels without altering its mRNA levels (Figure 3E&F), and specifically reduced the association of *Lsd1* mRNA with polyribosomes (Figure 3G&H). Altogether, these results indicated that the translation of *Lsd1* mRNA was specifically repressed by the R783H Trim71 mutant.

**Figure 3.**
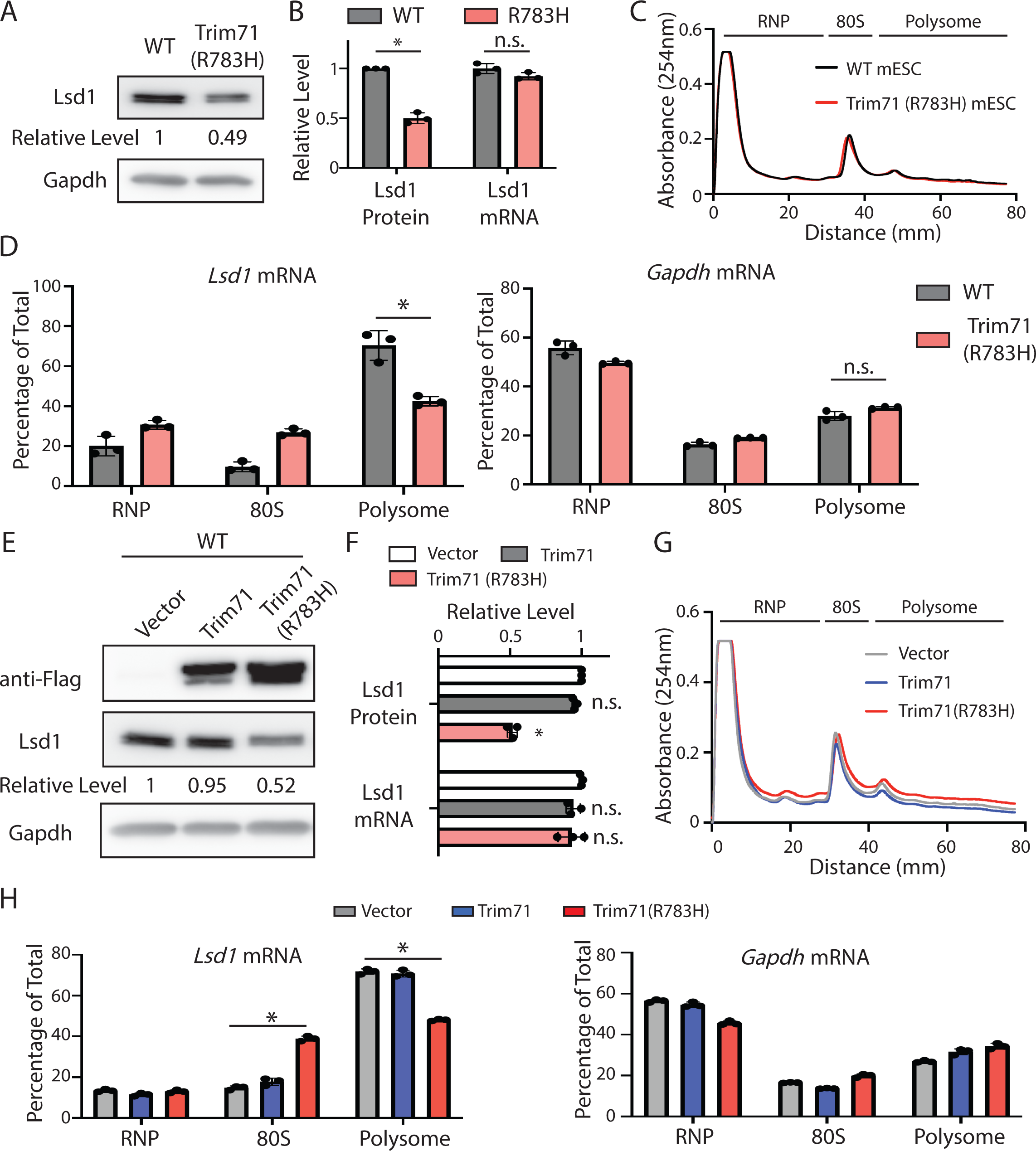
The Trim71 mutant R783H represses *Lsd1* mRNA translation in mESCs. A. Western blotting in the WT and the Trim71 (R783H) mESCs. B. Quantification of Lsd1 protein and mRNA levels. Gapdh and 18S rRNA were used for normalization in protein and mRNA quantifications, respectively. C. Polysome analysis in the WT and the Trim71 (R783H) mESCs. D. Quantification of the indicated mRNA distribution in the RNP, 80S, and polysome fractions from the WT and the Trim71 (R783H) mESCs. E. Western blotting in the WT mESCs expressing an empty vector, Flag-Trim71, Flag-Trim71(R783H). F. Quantification of Lsd1 protein and mRNA in the WT mESCs expressing an empty vector, Flag-Trim71, Flag-Trim71(R783H). G. Polysome analysis in the WT mESCs expressing an empty vector, Flag-Trim71, Flag-Trim71(R783H). H. Quantification of the indicated mRNA distribution in the RNP, 80S, and polysome fractions from the WT mESCs expressing an empty vector, Flag-Trim71, Flag- Trim71(R783H). The quantification results in B, D, F, and H represent the means (± SD) of three independent experiments. *p<0.05; and n.s. not significant (p>0.05) by the Student’s t-test. The following figure supplement is available for Figure 3: Figure supplement 1. Repression of *Lsd1* mRNA translation by the Trim71 mutant R783H is dependent on its binding to *Lsd1* mRNA. Figure supplement 2. The human TRIM71 mutant R796H represses LSD1. Figure 3 - source data Tiff files of raw gel images for Figure 3A, E, Figure 3-figure supplement 1A, Figure3- figure supplement 2.

This repression is dependent on the binding of the mutant Trim71 to the 3’UTR of *Lsd1* mRNA. Because in the Lsd1 CLIPΔ mESCs, where the interaction between *Lsd1* mRNA 3’UTR and the mutant Trim71 was abolished (see below), the mutant Trim71 failed to decrease both Lsd1 protein level and *Lsd1* mRNA’s association with polyribosomes in mESCs (Figure 3-figure supplement 1).

The sequence of the *Lsd1* mRNA 3’UTR is not conserved between mouse and human. However, similar stem-loop structures recognized by the mutant Trim71 (Figure 2C) are predicted *in silico* to be present in the 3’UTR of human *LSD1* mRNA, suggesting that the corresponding mutant human TRIM71 may also be able to repress *LSD1*. To test this, we expressed WT and the corresponding R796H mutant TRIM71 in NCCIT cells, which are human embryonal carcinoma cells. The human mutant TRIM71, but not WT TRIM71, reduced LSD1 protein levels without significantly changing *LSD1* mRNA levels (Figure 3-figure supplement 2), similar to the results in mESCs (Figure 3E&F), indicating the ability of the CH-causing Trim71 mutation to repress Lsd1 expression is conserved between mouse and human.

### Specific inhibition of *Lsd1* repression alleviates stem cell and neural differentiation defects

To evaluate the functional relevance of the mutant-Trim71-mediated translational repression of *Lsd1* mRNA to the differentiation defects of the mutant mESCs, we generated bi-allelic deletion of the mutant Trim71-binding region (∼60bp), defined from the CLIP-seq (Figure 2G), in the 3’UTR of *Lsd1* using genome editing (Figure 4A). We named this deletion as Lsd1 CLIPΔ. CLIP-qRT-PCR indicated that in the Lsd1 CLIPΔ mESCs the interaction between *Lsd1* mRNA and the mutant Trim71 was specifically disrupted. Because the mutant Trim71 did not bind *Lsd1* mRNA, but still specifically interacted with other target mRNAs, such as *Cdkn1a* mRNA and *Ddx6* mRNAs (Figure 4B). Thus, the Lsd1 CLIPΔ enabled us to specifically examine the functional significance of the mutant-Trim71:*Lsd1*-mRNA interaction at both molecular and cell function levels.

**Figure 4.**
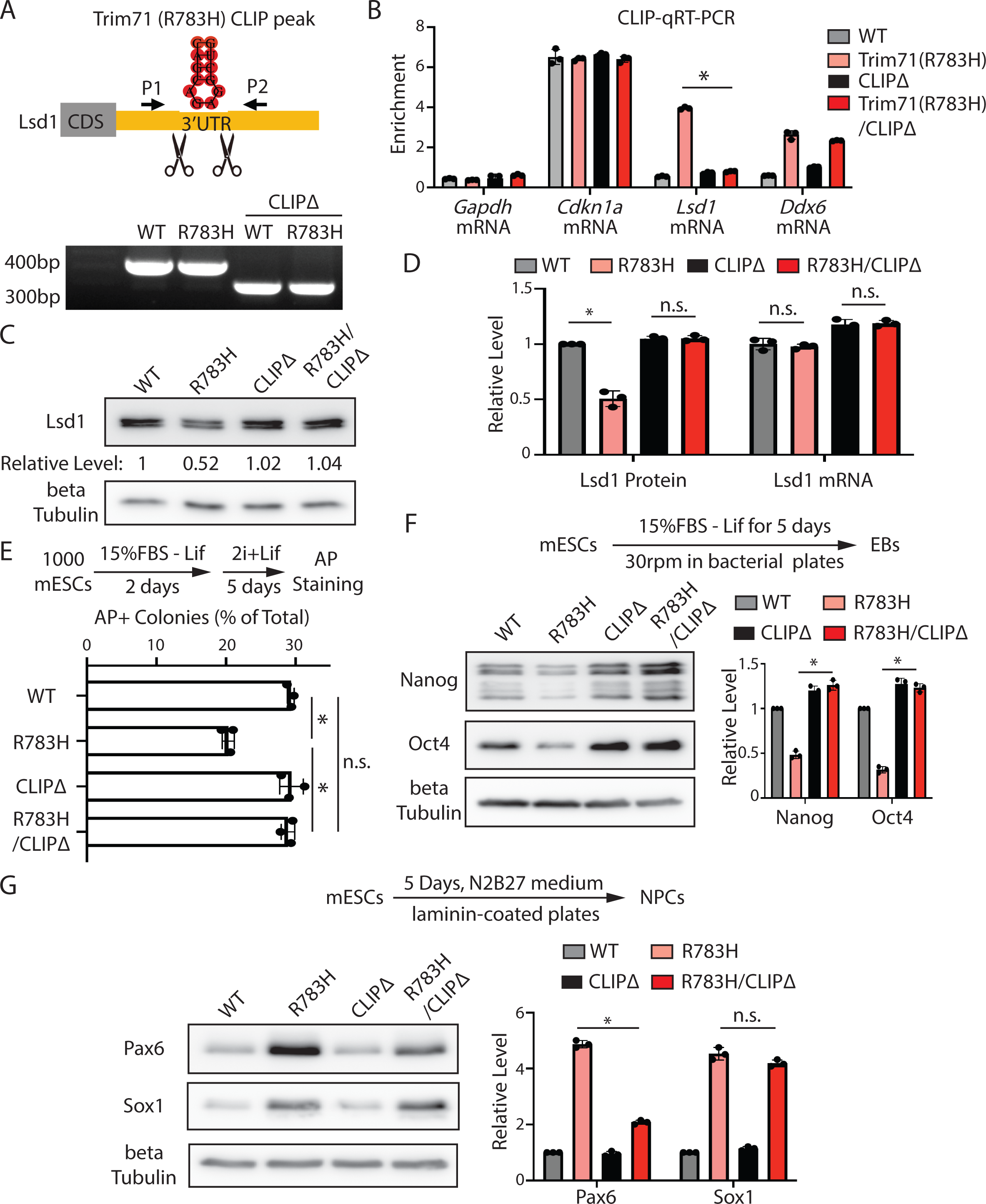
Specific inhibition of the interaction between the Trim71 mutant R783H and *Lsd1* mRNA alleviates the stem cell and neural differentiation defects in the Trim71(R783H) mESCs. A. Deletion of the Trim71 mutant R783H binding site in *Lsd1* mRNA’s 3’UTR. B. CLIP-RIP followed by qRT-PCR to examine mRNAs associated with the Trim71 and the Trim71 mutant R783H in the WT, Trim71(R783H), CLIPΔ, and Trim71(R783H)/CLIPΔ mESCs. The mRNA signals from the E14 mESCs were set as 1 for relative comparison. C. Western blotting in the WT, Trim71(R783H), CLIPΔ, and Trim71(R783H)/CLIPΔ mESCs. D. Quantification of Lsd1 protein and mRNA in the WT, Trim71(R783H), CLIPΔ, and Trim71(R783H)/CLIPΔ mESCs. Beta-Tubulin and 18S rRNA were used for normalization in the protein and mRNA quantification, respectively. E. Exit pluripotency assay for mESCs. F. Representative Western blotting and quantification of pluripotency factors during EB formation. G. Representative Western blotting and quantification of neural lineage markers during mESCs neural differentiation. The results from B, D, F, and G represent the means (± SD) of three independent experiments. In E, the colony morphology and AP intensity were evaluated through microscopy. 100-200 colonies were examined each time to determine the percentage of undifferentiated colonies. *p<0.05; and n.s. not significant (p>0.05) by the Student’s t- test. The following figure supplement is available for Figure 4: Figure supplement 1. The R783H mutation in the Lsd1 CLIPΔ background does not alter the polysome association of Lsd1 mRNA. Figure 4 - source data Tiff files of raw gel images for Figure 4A, C, F, and G.

At the molecular level, the Lsd1 CLIPΔ increased Lsd1 protein in the mutant Trim71 mESCs to a level similar to that in WT mESCs (Figure 4C&D). *Lsd1* mRNA, however, was not significantly increased (Figure 4D), further confirming the translational repression mediated by the R783H Trim71 mutant. Notably, unlike the observations in R783H Trim71 mutant mESCs, the Lsd1 CLIPΔ did not increase Lsd1 protein levels in WT mESCs (Figure 4C), indicating that the Lsd1 CLIPΔ sequence in the 3’UTR of *Lsd1* mRNA does not regulate Lsd1 production *in cis*, but controls *Lsd1* mRNA translation through interacting with the mutant Trim71. Moreover, in the Lsd1 CLIPΔ background, the R783H mutation did not alter the polysome association of *Lsd1* mRNA (Figure 4- figure supplement 1), which is different from the results in the WT background (Figure 3C&D), indicating that the translation repression requires the binding of the R783H Trim71 mutant to *Lsd1* mRNA. These findings, combined with the R783H Trim71 mutant CLIP-seq results, revealed that the CH-causing mutation significantly alters, but does not abolish, RNA target recognition by Trim71.

At the cell function level, the Lsd1 CLIPΔ abolished the stem cell differentiation defects of the R783H Trim71 mutant mESCs, as indicated by both the exit pluripotency assay (Figure 4E) and the expression level of pluripotency markers during EB formation (Figure 4F). Moreover, during neural differentiation, the Lsd1 CLIPΔ alleviated the accelerated neural differentiation in the mutant Trim71 mESCs. This was manifested as a decrease in Pax6 when the Lsd1 CLIPΔ was introduced in the mutant mESCs (Figure 4G). However, no significant changes of Sox1 were observed (Figure 4G). This is possibly due to additional functional targets of the R783H Trim71 mutant during neural differentiation. Nevertheless, these results collectively indicate that repression of *Lsd1* mRNA translation by the R783H Trim71 mutant is required for the stem cell and neural differentiation defects in the R783H Trim71 mESCs.

### Increasing Lsd1 alleviates the stem cell and neural differentiation defects

To evaluate the functional significance of Lsd1 in the differentiation defects seen in the CH-causing R783H mutation, we asked whether increasing Lsd1 protein levels could mitigate the phenotypes of the mutant mESCs. For this purpose, we generated stable mESC lines in which the expression of an Lsd1-GFP fusion protein can be induced by doxycycline (dox) in a dosage-dependent manner (Figure 5A). The GFP fusion enabled us to discriminate exogenous Lsd1 from endogenous Lsd1. Modulation of dox levels revealed that a ∼25% increase in Lsd1 protein levels over endogenous levels alleviated the differentiation defects seen in the mutant mESCs, as revealed by both the exit pluripotency assay (Figure 5B) and the expression level of pluripotency markers during differentiation (Figure 5C). This alleviation is specific to the mutant mESCs, as expression of exogenous Lsd1 at a similarly increased level did not cause phenotypical changes in WT mESCs (Figure 5A-C). Moreover, this alleviation requires the demethylase activity of Lsd1, because expressing a demethylase catalytic mutant of Lsd1 failed to mitigate the differentiation defects of the mutant mESCs (Figure 5-figure supplement 1). During neural differentiation, similar to the result from the Lsd1 CLIPΔ approach (Figure 4G), although not decreasing Sox1 to the normal level, the slightly (∼25%) increased Lsd1 reduced the overexpressed Pax6 in the mutant mESCs (Figure 5D), indicating that increasing Lsd1 mitigates the neural differentiation defects in the mESCs with the CH-causing mutation. Thus, decreased Lsd1 protein levels in R783H Trim71 mutant mESCs plays a critical role in the observed stem cell and neural differentiation defects.

**Figure 5.**
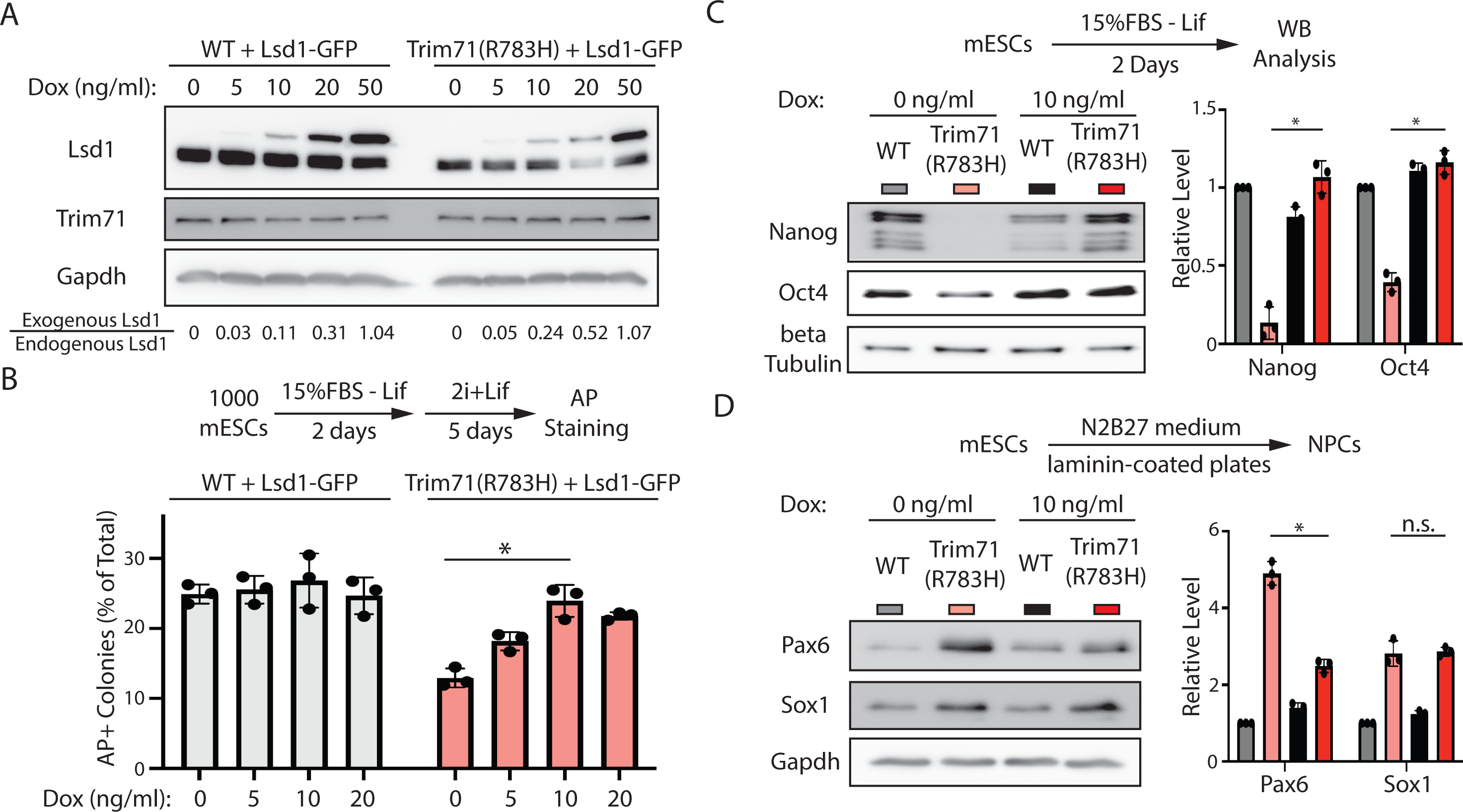
A slight increase of Lsd1 alleviates the stem cell and neural differentiation defects in the Trim71(R783H) mESCs. A. Western blotting in the WT and the Trim71(R783H) mESCs with dox-inducible expression of Lsd1-GFP. B. Exit pluripotency assay for mESCs. C. Representative Western blotting and quantification of pluripotency factors during the monolayer differentiation of mESCs. D. Representative Western blotting and quantification of neural lineage markers during mESCs neural differentiation. In B, the colony morphology and AP intensity were evaluated through microscopy. 100- 200 colonies were examined each time to determine the percentage of undifferentiated colonies. The quantification results from C and D represent the means (± SD) of three independent experiments. *p<0.05; and n.s. not significant (p>0.05) by the Student’s t- test. The following figure supplement is available for Figure 5: Figure supplement 1. The demethylase catalytic mutant Lsd1 fails to alleviate the stem cell differentiation defects in the Trim71(R783H) mESCs. Figure 5 - source data Tiff files of raw gel images for Figure 5A, C, D, and Figure 5-figure supplement 1A, C.

### Lsd1 plays an important role in the differentiation defects in mESCs with monoallelic R783H mutation on Trim71

A caveat of the above results is that although the bi-allelic R783H Trim71 mutation in mESCs provides critical functional and mechanistic insights into the mutant Trim71, it is the monoallelic R783H mutation that causes CH. To evaluate whether the mechanistic insights we obtained using bi-allelic R783H mESCs are relevant to the pathogenesis of the disease, we examined mESCs with a monoallelic R783H mutation in Trim71 (R783H/+) (Figure 1-figure supplement 1B), which mimics the genetic setting of CH. The R783H/+ mESCs displayed similar stem cell and neural differentiation defects as the bi-allelic R783H mESCs, as indicated by decreased Lsd1 protein levels (Figure 6A), rapid exit from pluripotency (Figure 6B&C), and accelerated neural differentiation (Figure 6D).

**Figure 6.**
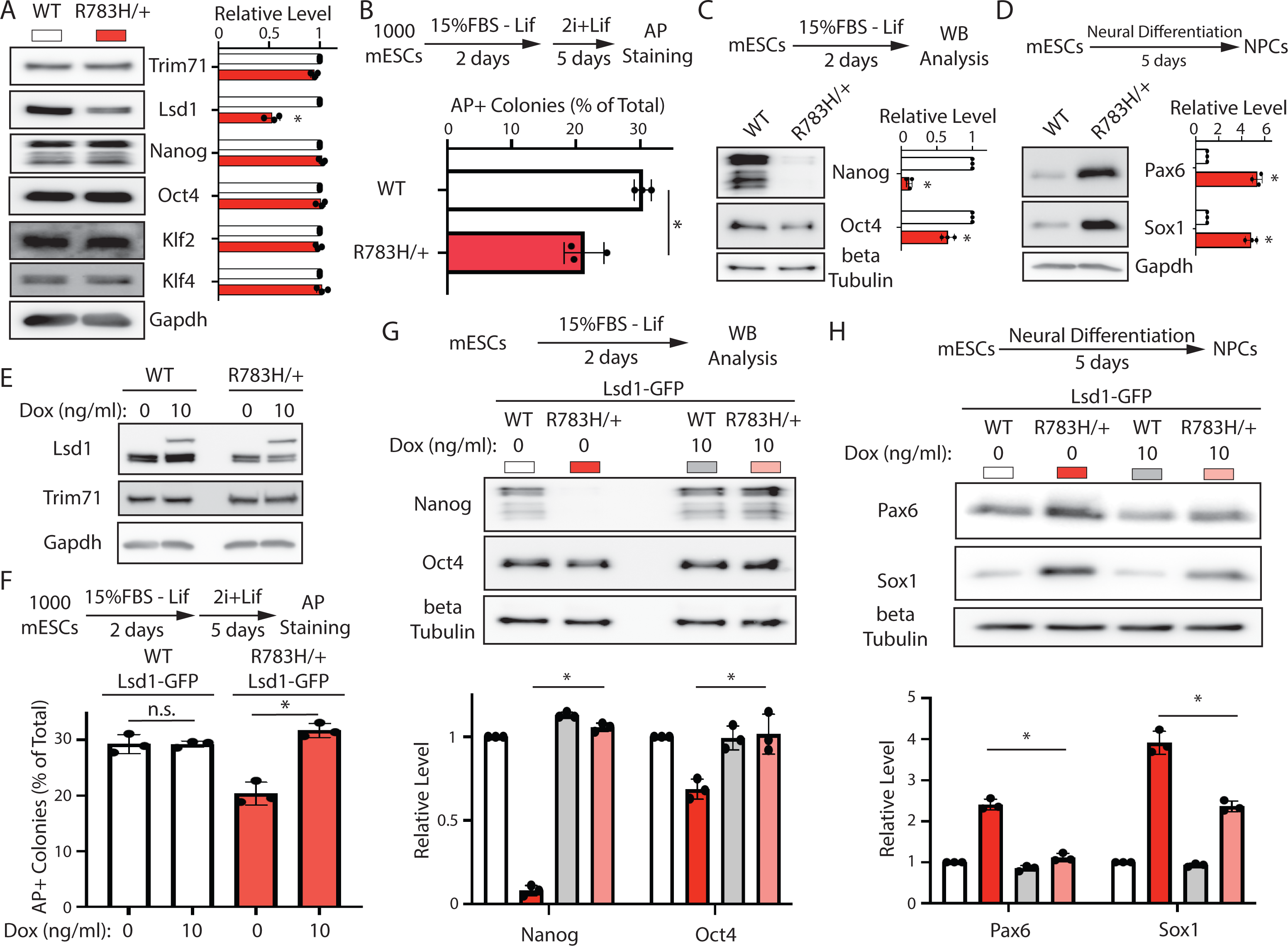
Monoallelic R783H mutation on Trim71 results in stem cell and neural differentiation defects in a Lsd1 dependent manner. A. Representative Western blotting and quantification in the WT and the R783H/+ mESCs. B. Exit pluripotency assay for the WT and the R783H/+ mESCs. C. Representative Western blotting and quantification of pluripotency factors in differentiating mESCs. D. Representative Western blotting and quantification of neural lineage markers during mESCs neural differentiation. E. Western blotting in the WT and the R783H/+ mESCs with dox-inducible expression of Lsd1-GFP. F. Exit pluripotency assay for the WT and the R783H/+ mESCs with dox-inducible expression of Lsd1-GFP. G. Representative Western blotting and quantification of pluripotency factors during the differentiation of the WT and the R783H/+ mESCs with dox-inducible expression of Lsd1-GFP. H. Representative Western blotting and quantification of neural lineage markers during the neural differentiation of the WT and the R783H/+ mESCs with dox- inducible expression of Lsd1-GFP. In B and F, the colony morphology and AP intensity were evaluated through microscopy. 100-200 colonies were examined each time to determine the percentage of undifferentiated colonies. The quantification results from A-D and F-H represent the means (± SD) of three independent experiments. *p<0.05; and n.s. not significant (p>0.05) by the Student’s t-test. Figure 6 - source data Tiff files of raw gel images for Figure 6A, C, D, E, G, and H. Supplemental file 1. CLIP-seq peaks from the WT Trim71 and the R783H Trim71 mutant in mESCs. Supplemental file 2. Antibodies, plasmids, and oligonucleotides used in this study.

To determine whether increasing Lsd1 level reduce these stem cell and neural differentiation defects in mESCs with the monoallelic mutation, we used dox-inducible expression of Lsd1-GFP in the R783H/+ mESCs (Figure 6E). Expression of exogenous Lsd1-GFP in R783H/+ mESCs alleviated the defects in stem cell differentiation (Figure 6F&G) and neural lineage commitment (Figure 6H). Thus, in the CH-mimic setting, a slight increase in Lsd1 protein levels can also mitigate the stem cell differentiation defects caused by the mutant Trim71.

## Discussion

Here, we show that, in mouse embryonic stem cells, the CH-causing R783H Trim71 gain-of-function mutation resulted in defects in stem cell differentiation and neural lineage commitment, both in the mono-allelic condition, as found in human patients, and the bi-allelic condition. Mechanistically, the R783H Trim71 mutation significantly changes the mRNA substrates to which Trim71 binds and potentially regulates in mESCs. Among the newly acquired mRNA substrates, we determined that the mutant Trim71 represses *Lsd1* mRNA translation. Specific inhibition of *Lsd1* translational repression or mild overexpression of Lsd1, both of which increase Lsd1 protein levels, alleviate the differentiation defects in mESCs expressing the CH-causing R783H Trim71 mutation. These results provide mechanistic insights into the pathogenesis of CH mediated by the R783H mutation in Trim71 and argue that Lsd1 can be a potential therapeutic target for CH.

### Trim71 target recognition

Our results reveal that the R783H mutation in Trim71 significantly changes its binding specificity for target mRNAs (Figure 2D). Given that both WT Trim71 and R783H Trim71 interact with mRNAs that share similar predicted secondary structure motifs (Figure 2C), the molecular basis for their differing target specificities is unclear. Trim71, a highly conserved RBP, binds target RNAs through its NHL domain. Structural and *in vitro* binding studies indicate that the binding specificity of the NHL domain is determined by the shape of an RNA stem-loop structure and not by the primary sequence motifs (Kumari *et al*., 2018). Based on a crystal structure of the NHL domain from *D*. *melanogaster* Brat (Loedige *et al*, 2015), a close homolog of Trim71, the R796H mutation (R783H in mouse) is predicted to alter the interaction between Trim71 and RNA’s phosphate backbone (Furey *et al*., 2018). We speculate that the point mutation impacts RNA structural shape recognition, altering target mRNA binding. Consistent with this notion, comparison of the CLIP-seq datasets revealed that the major difference between WT and mutant Trim71 is a decreased stringency in the stem region of the predicted stem-loop/hairpin structure enriched in the mutant-Trim71-binding regions (Figure 2C).

However, the enriched structural motifs identified using CLIP-seq are likely necessary, but not sufficient, for the binding. Because similar structural motifs can be predicted *in silico* outside Trim71-binding regions defined by CLIP-seq in target mRNAs and in non-target mRNAs. This implies that besides the structural motifs, additional features are involved in Trim71’s target recognition. Furthermore, it is unclear whether and when the *in silico* predicted secondary structures are formed *in vivo*. Unlike *in vitro* folding, formation of RNA structures *in vivo* is constrained by contexts (e.g., RBPs in the neighboring region, etc.). Thus, future structural studies on Trim71 combined with *in vivo* probing of RNA structures will reveal how Trim71 specifically recognizes its targets and how mutations alter this process.

### Characterization of functional RBP:mRNA interactions

CLIP is widely used in identifying *in vivo* RBP:RNA interactions (Hafner *et al*, 2021; Lee & Ule, 2018); and, when combined with high throughput sequencing, this method has revealed transcriptome-wide binding sites and the corresponding target genes for many RBPs (Van Nostrand *et al*, 2020). However, opportunistic or non-productive interactions can complicate identifying functional targets. While loss-of-function (knockout/knockdown) and gain-of-function (overexpression) approaches can determine RBPs’ functions, these methods, however, provide limited insights into the significance of specific RBP:RNA interactions. Because an RBP usually binds and potentially regulates numerous RNAs, and knockout or overexpression of an RBP can lead to alteration in many RBP:RNA interactions, making assigning any phenotypic changes to specific RBP:RNA interactions challenging.

Here we specifically disrupted the interaction between the R783H Trim71 mutant and *Lsd1* mRNA, by deleting the 3’UTR binding site identified using CLIP-seq (Figure 4A). This approach does not abolish the interactions between the mutant Trim71 and its other target mRNAs (Figure 4B), thereby we could specifically evaluate the role of this interaction plays in the R783H Trim71 mediated mESC differentiation defects. We believe similar approaches will reveal many more functional RBP:mRNA interactions in normal and pathological processes.

### Lsd1 and neural differentiation

Lsd1 is a conserved histone lysine-specific demethylase with critical functions in stem cell biology (Adamo *et al*, 2011; Whyte *et al*., 2012). Besides histone, Lsd1 can also modulate the methylation status of other proteins (e.g., p53) (Perillo *et al*, 2020).

Previous studies indicated that the elimination of Lsd1 through proteasome-mediated degradation promotes mESC differentiation toward neural lineage (Han *et al*., 2014). Here we found that the CH-causing Trim71 mutant (R783H) binds to *Lsd1* mRNA and repress its translation, leading to accelerated stem cell differentiation and premature neural lineage commitment.

When regulating histones in chromatin, Lsd1 can both activate and repress gene expression through association with different histone modification complexes (Kozub *et al*, 2017). We found that the ∼50% reduction in Lsd1 protein levels in R783H Trim71 mESCs, mediated by translational repression, results in differentiation defects (Figure 4), while a ∼25% increase in Lsd1 protein levels in the same R783H Trim71 mESCs alleviates these defects (Figure 5). These observations argue that genes controlled by weak/dynamic Lsd1 binding may mediate the differentiation defects in the mutant mESCs. Because weak/dynamic interactions are more sensitive to concentration fluctuations than strong/steady interactions. Numerous Lsd1 target genes have been identified using ChIPs (chromatin immunoprecipitations), however the formaldehyde crosslinking step makes quantitatively discriminating weak versus strong chromatin:protein interactions challenging. Thus, non-crosslinking approaches, such as CUT&Tag (Kaya-Okur *et al*, 2019), may be more suitable for identifying those weak/dynamic Lsd1 chromatin targets altered in the mutant mESCs. It is also possible that the decreased Lsd1 in the mutant mESCs may change the methylation status, thereby modulating the activity/function, of non-histone proteins. Characterizing such potential functional targets of Lsd1 may reveal novel regulators of neural differentiation.

Finally, it is important to mention that although we identified Lsd1 as a critical functional target of the R783H Trim71 mutant, it is likely not the only functional target, because the altered Lsd1 protein levels can only partially explain the neural differentiation defects. Additional functional target(s) of Trim71 with the CH-causing mutation may also contribute to these cellular defects.

## Materials and Methods

All the antibodies, oligonucleotides, and plasmids used in this study are listed in Supplementary file 2.

### Cell Culture

All the mouse ESCs used in this study are derived from ES-E14TG2a (ATCC CRL- 1821). mESCs were cultured on 0.5% gelatin-coated tissue cultured plates, in either 15%FBS + Lif medium or 2i+Lif medium (Mulas *et al*., 2019). Human cell line NCCIT (ATCC, CRL-2073) was maintained in RPMI-1640 medium supplemented with 10% FBS. All cells were incubated at 37 °C with 5% CO_2_.

### CRISPR/Cas9-mediated Genome Editing in mESCs

To generate the FLAG-Trim71R783H mESCs and FLAG-Trim71R783H+/- mESCs, FLAG-Trim71 mESCs were co-transfected with 2 µg of pWH464 (pSpCas9(BB)−2A- GFP (pX458)) carrying the targeting sgRNA (oWH4229) and 1 µg of donor oligo (oWH4189) using Fugene6. To generate Lsd1 CLIPΔ cells, 2 µg of pWH464 expressing a pair of sgRNAs targeting the indicated region were transfected into the mESCs.

Transfected cells were single-cell sorted to 96-well plates. Colonies were then picked and expanded for validation by genotyping PCR followed by sequencing and Western blot analysis.

### Generation of Stable Cell Lines

Stable cell lines expressing doxycycline-inducible mouse FLAG-Trim71, mouse FLAG- Trim71R783H, mouse LSD1-GFP, human 3xHA-Trim71, or human 3xHA-Trim71R796H were generated using a PiggyBac transposon-based expression system. Briefly, cells were co-transfected with indicated plasmids and PiggyBac transposase (pWH252).

After 48hrs, cells were selected with 1 µg/ml puromycin for 4 days.

### RNA extraction and RT-qPCR

RNA was extracted from cells using RNA reagent and treated with DNase1 to remove contaminating DNA. cDNA was synthesized using random hexamers and Superscript2 reverse transcriptase (Invitrogen) according to manufacture instructions. qPCR was performed in triplicate for each sample using the SsoAdvanced Universal SYBR Green Supermix (Bio-Rad) and a CFX96^TM^ real-time PCR detection system (Bio-Rad).

### Western Blotting

Proteins were harvested in RIPA buffer (10 mM Tris-HCl pH 8.0, 140 mM NaCl, 1 mM EDTA, 1% Triton X-100, 0.5 mM EGTA, 0.1% SDS, 0.1% sodium deoxycholate, and protease inhibitor cocktail) and quantified with BCA assay kit. Protein samples were resolved by SDS-PAGE and then transferred to PVDF membranes. Western blotting was performed using a BlotCycler (Precision Biosystems) with the indicated antibodies. Signals were developed with the Western ECL substrate (Bio-Rad) and detected with an ImageQuant LAS 500 instrument (GE Healthcare).

### Colony formation assay and exit from pluripotency assay

For colony formation assay, 500 cells per well were plated on a 12-well plate in 15% FBS +Lif medium for 6 days. For exit from pluripotency assay, 1000 cells per well were plated on 6-well plate in differentiation media (15%FBS – Lif) for 2 days, then cultured in 2i+Lif medium for another 5 days. Colonies were stained using an Alkaline Phosphatase Assay Kit (System Biosciences) and evaluated under an Olympus CK2 microscope.

### Embryoid body formation and monolayer differentiation

For embryoid body formation assay, 3 × 10^6^ mES cells were seeded into 10 cm bacterial grade Petri dish in 10 ml differentiation medium (DMEM/F12 supplemented with 15% FBS, 1 × penicillin/streptomycin, 0.1 mM Non-Essential Amino Acids, 2 mM L-glutamine, and 50 uM 2-mercaptoethanol), and maintained on a horizontal rotator with a rotating speed of 30 rpm. The medium was changed at day 3 and the resultant EBs were harvested at day 5. For monolayer differentiation, 1× 10^4^ mES cells per well were seeded into gelatin-coated 6-well plate in 2ml differentiation medium for 2 days.

### Neural cell differentiation

mESCs were dissociated and seeded onto 10 ug/ml laminin-coated 6-well plate at a density of 1 x 10^4^ cells/cm^2^ in N2B27 medium (Mulas *et al*., 2019). The medium was changed on day 2 and every day thereafter. Cells were harvested at the indicated time points.

### Polysome analysis

Polysome analysis was performed using the methods described previously (Zhang *et al*, 2017). Briefly, mESCs were lysed in the polysome lysis buffer (10 mM Tris-HCl pH 7.4, 12 mM MgCl2, 100 mM KCl, 1% Tween-20, and 100 mg/mL cycloheximide). Then 5 OD260 cell lysate was loaded onto a 5%–50% (w/v) linear sucrose-density gradient, followed by centrifugation at 39,000 rpm in a Beckman SW-41Ti rotor for 2 hr at 4°C. The gradient was fractionated using a Gradient Station (BioComp) coupled with an ultraviolet 254nm detector (Bio-Rad EM-1).

### CLIP-seq and peak calling, and RNA-seq

CLIP-seq was performed using the method described in the previous paper. The peak calling was performed using the pipeline described previously (Chen *et al*, 2019; Liu *et al*., 2021a). The CLIP-seq and RNA-seq datasets generated during this study are available at GEO: GSE183715 (reviewer access token: sfareaaynvuxlsz) and GSE196017 (reviewer access token: mzcxoiskjxepvex), respectively.

## Acknowledgments

This work is supported by Mayo Foundation for Medical Education and Research. We thank Drs. J. Alvarez-Dominguez and G. Riddihough for critical comments.

## Author Contributions

W.H. conceived the project and supervised the study. Q.L., M.K.N., R.M.P., K.R.M., and W.H. performed experiments and interpreted the data. X.C. and S.Z. performed the computational analysis. W.H. wrote the manuscript with inputs from all the authors.

---

**Figure 1-figure supplement 1.**
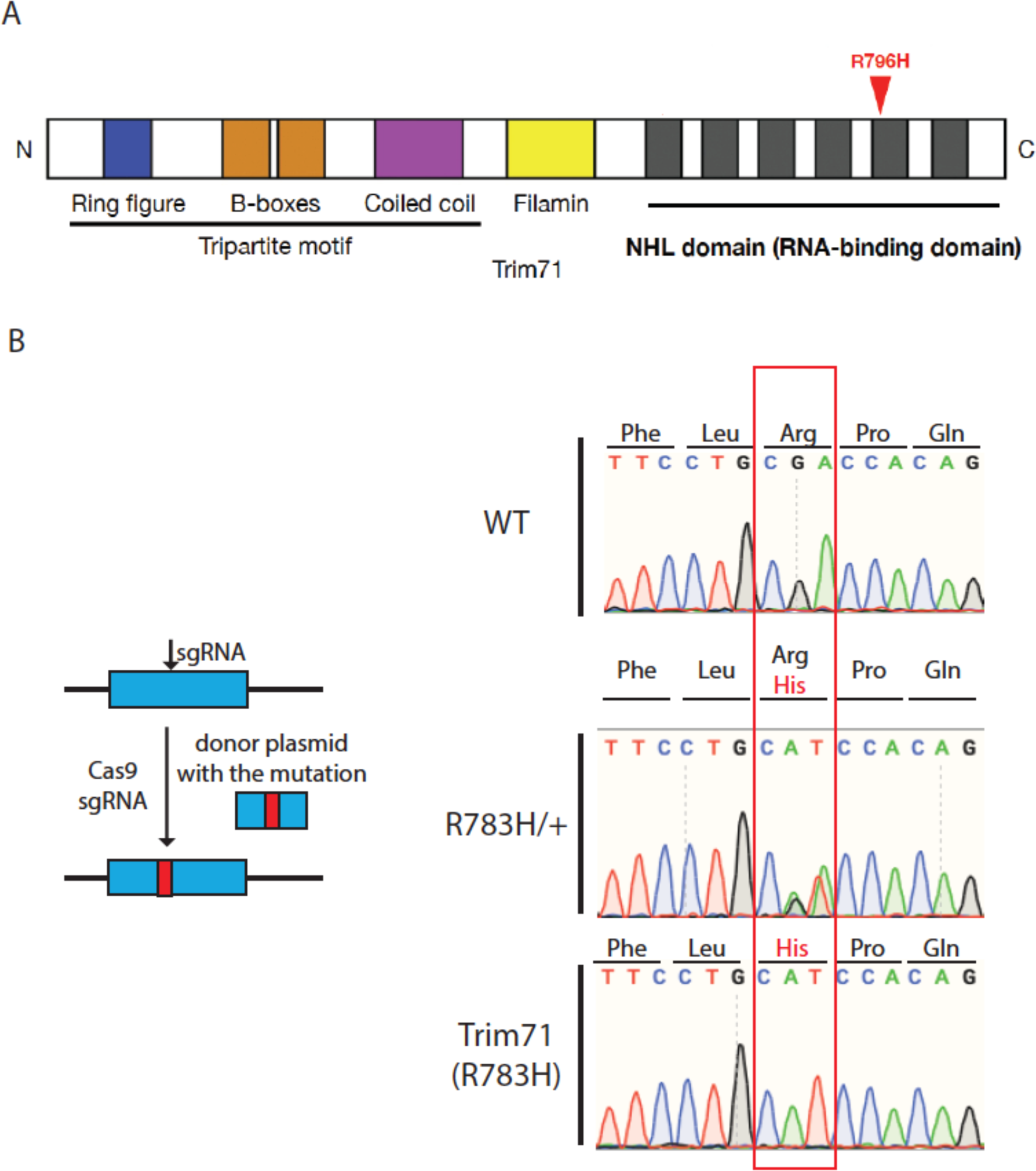
Generation of the Trim71 R783H monoallelic and bi-allelic mESCs. A. The location of R796H mutation on human Trim71 protein. The cartoon of Trim71 domain is adapted from Furey et al., 2018. B. Work flow of generating the R783H mutation in mESCs and sanger sequencing results verifying the mutation in mESCs.

**Figure 1-figure supplement 2.**
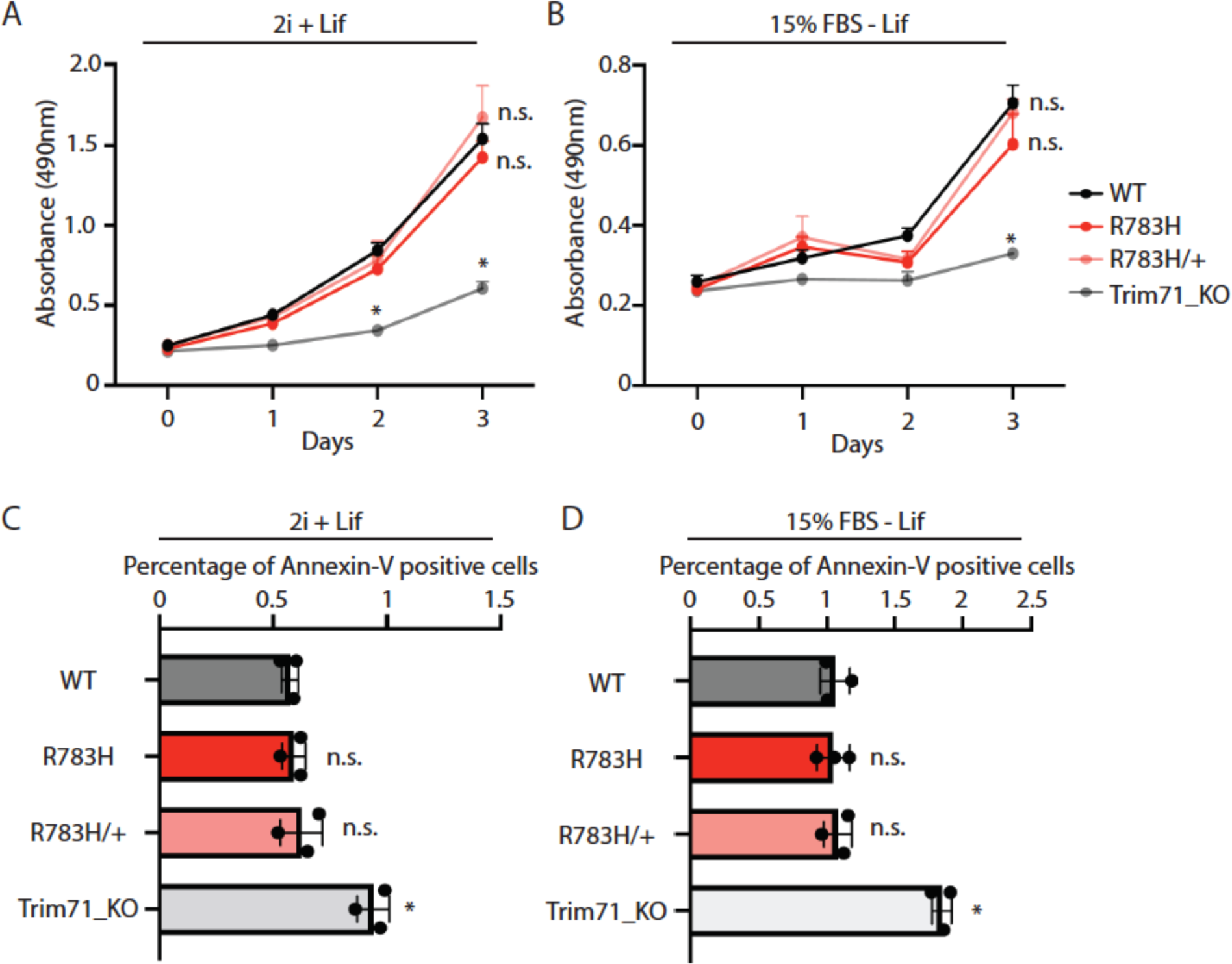
The R783H mutation in Trim71 does not change the proliferation and apoptosis of mESCs. A. Proliferation of mESCs under the stemness condition. The cells were cultured in the 2i+Lif medium, and the cell proliferation was monitored by the CellTiter 96 AQueous One Solution Cell Proliferation Assay (Promega). B. Proliferation of mESCs under the differentiating condition. The cells were cultured in the 15%FBS - Lif medium, and the cell proliferation was monitored by the CellTiter 96 AQueous One Solution Cell Proliferation Assay (Promega). C. Apoptosis of mESCs under the stemness condition. The cells were cultured in the 2i+Lif medium, and the cellular apoptotic state was monitored by annexin-V and PI staining followed by flow cytometry analysis. D. Apoptosis of mESCs under the differentiating condition. The cells were cultured in the 2i+Lif medium, and the cellular apoptotic state was monitored by annexin-V and PI staining followed by flow cytometry analysis. The results represent the means (± SD) of three independent experiments. * p < 0.05, n.s. not significant (p>0.05) by the Student’s t-test.

**Figure 1-figure supplement 3.**
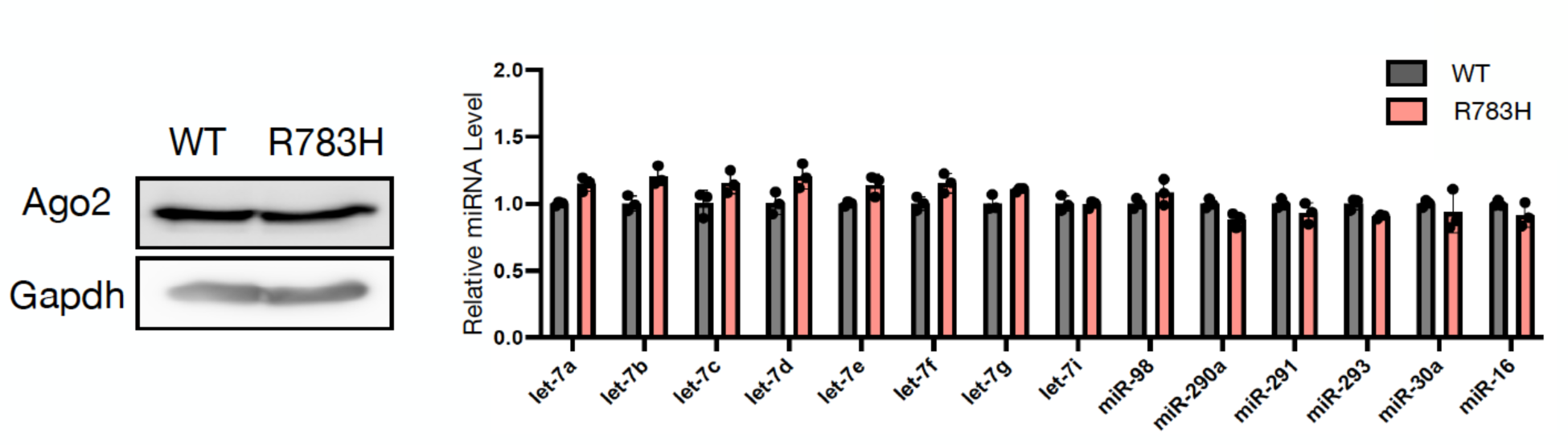
The R783H mutation in Trim71 does not alter the microRNA pathway in mESCs. The left panel shows a representative Western blot of Ago2 and Gapdh in the WT and R783H mESCs. The right panel shows the expression levels of a group of microRNAs in the WT and R783H mESCs. The microRNA levels were determined by qRT-PCR using U6 RNA levels for normalization. The qRT-PCR results represent the means (± SD) of three independent experiments, and none of the examined microRNAs have significantly different (p < 0.05) expression levels in the R783H mutant mESCs as determined by the Student’s t-test.

**Figure 1-figure supplement 4.**
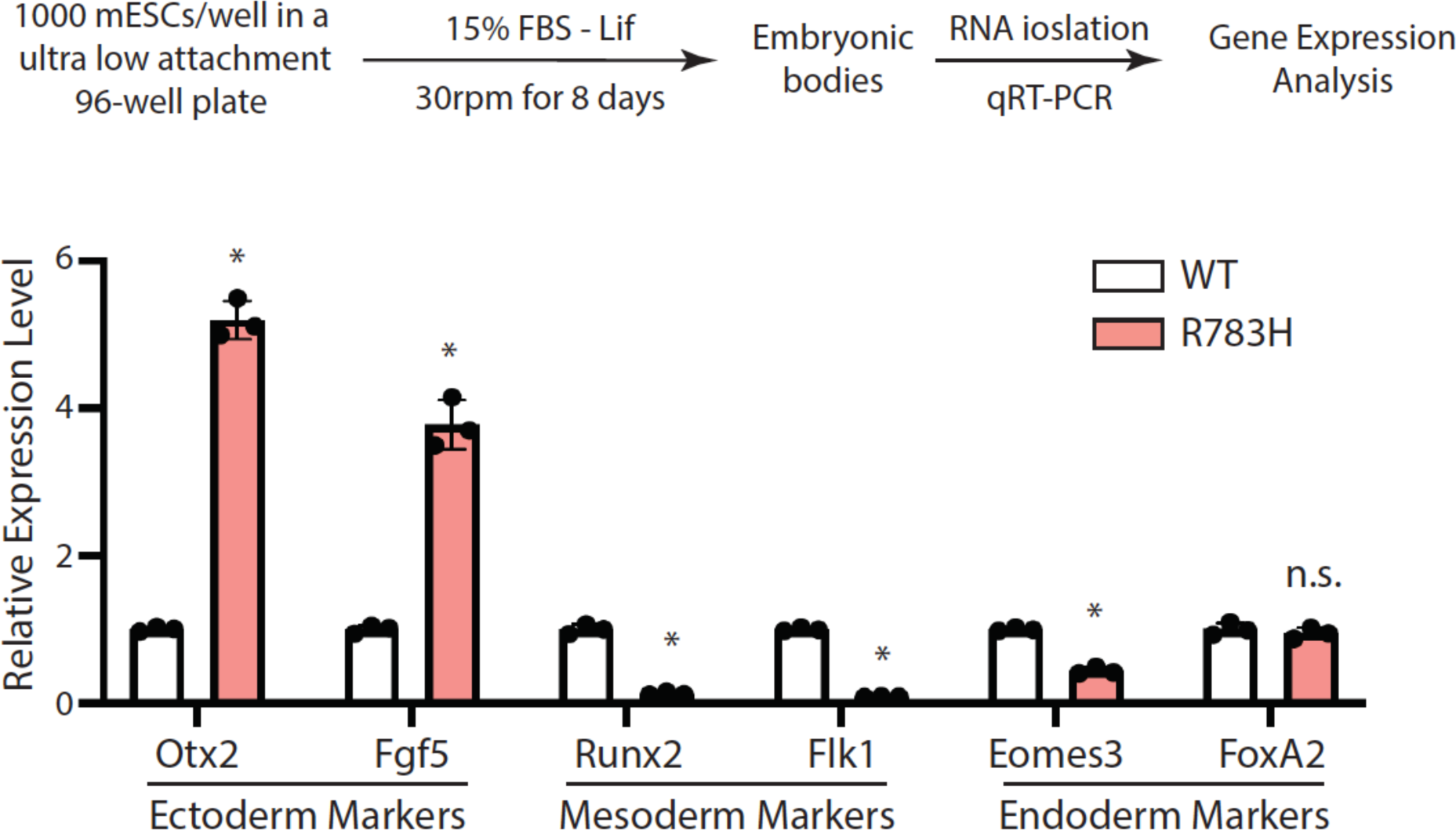
The R783H mESCs are more prone differentiate into the ectoderm lineage during EB formation. 18S rRNA was used for the normalization in the gene quantification by qRT-PCR. The qRT-PCR results represent the means (± SD) of three independent experiments. * p < 0.05, n.s. not significant (p>0.05) by the Student’s t-test.

**Figure 2-figure supplement 1.**
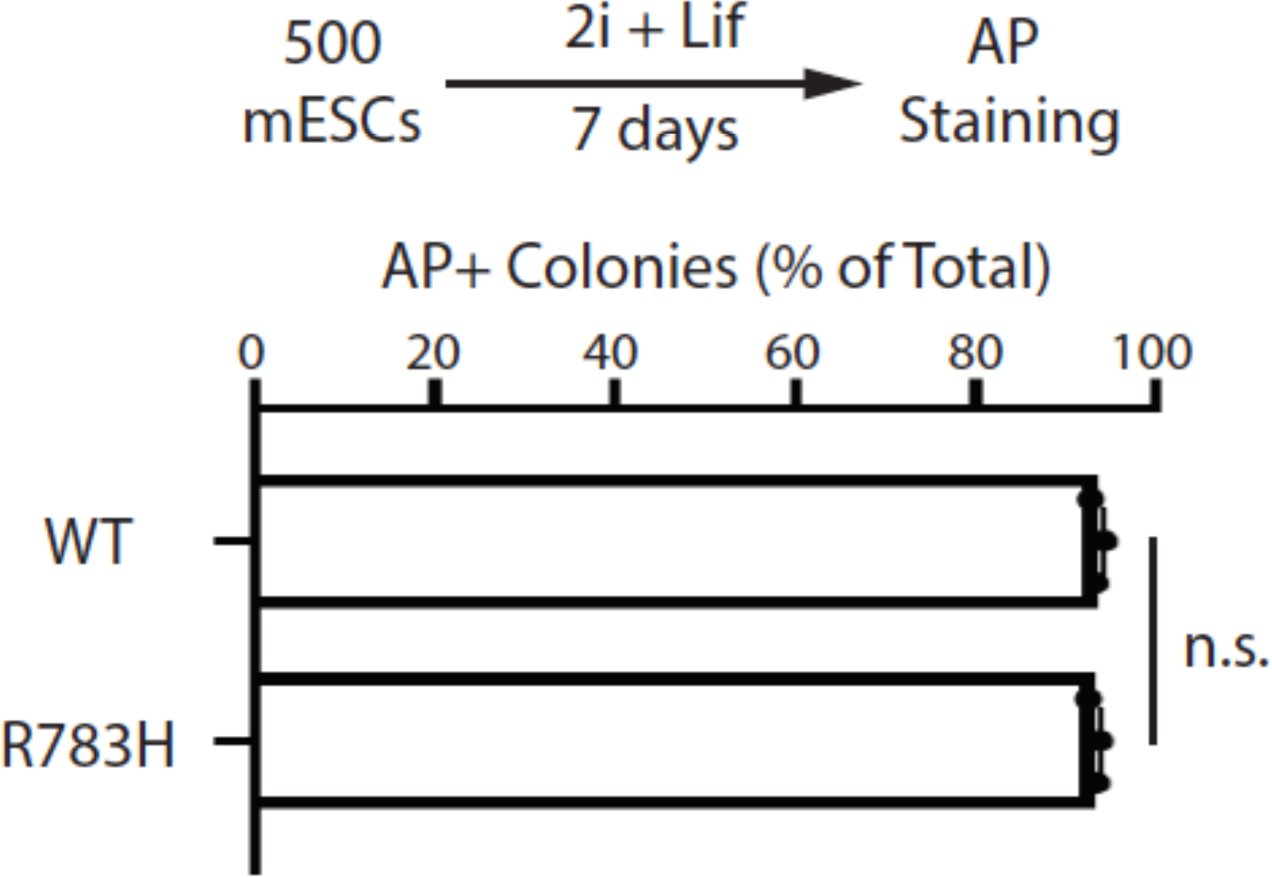
Colony formation assay for the WT and the R783H mESCs grown in the 2i+lif medium. The results represent the means (± SD) of three independent experiments. The colony morphology and AP intensity were evaluated through microscopy. 100-200 colonies were examined each time to determine the percentage of undifferentiated colonies. n.s. not significant (p>0.05) by the Student’s t-test.

**Figure 2-figure supplement 2.**
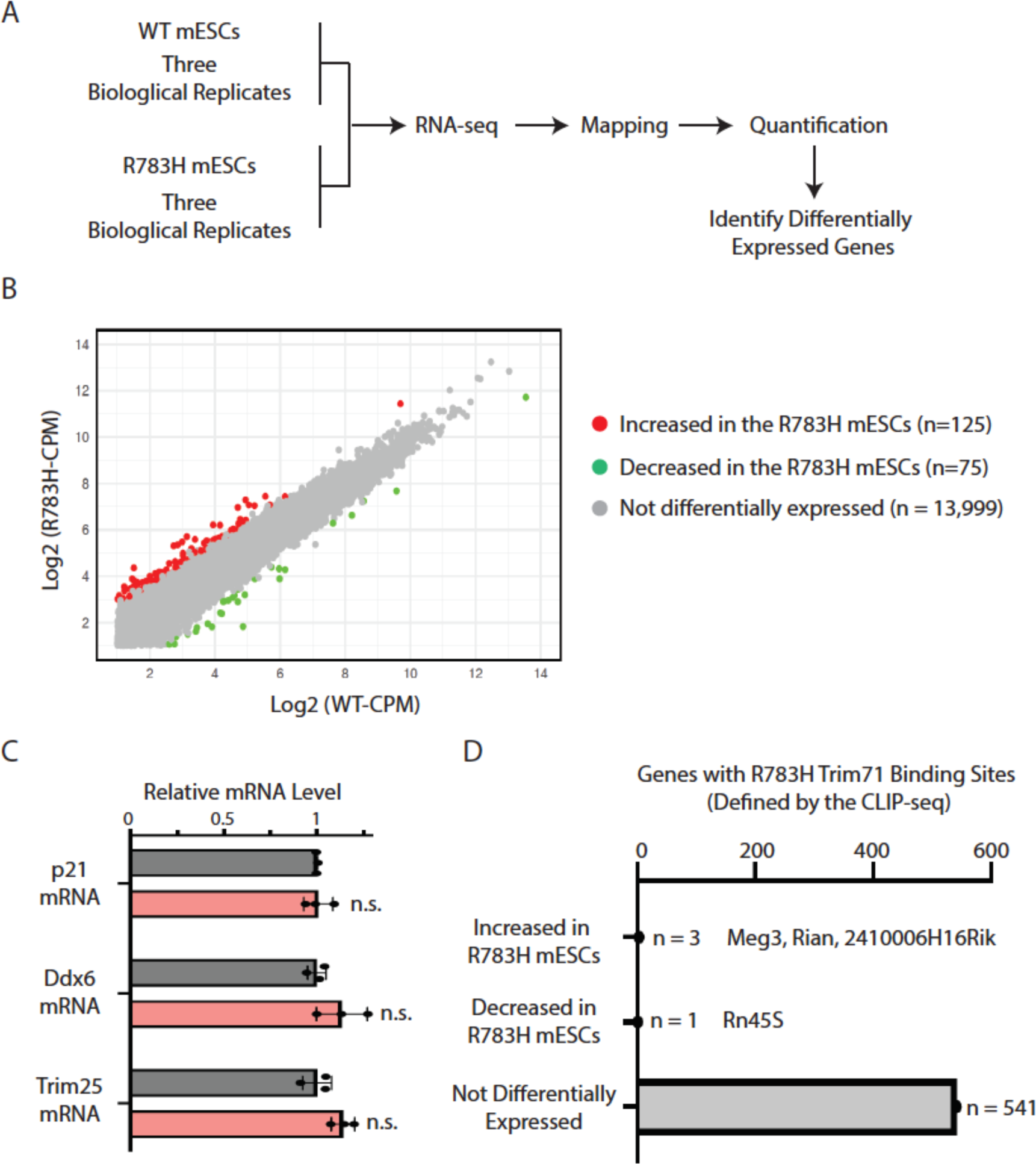
RNA-seq analysis of the WT and R783H mESCs. A. Work flow of the RNA-seq analysis. B. Differentially expressed genes in the WT and R783H mESCs. The differentially expressed genes were identified by the edgeR package. CPM: counts per million reads. The expression level of each gene is the average of the three biological replicates. C. qRT-PCR verification on several target mRNAs of the R783H Trim71 mutant. 18S rRNA was used for normalization. The results represent the means (± SD) of three independent experiments. n.s. not significant (p>0.05) by the Student’s t- test. D. Distribution of the target RNAs of the R783H Trim71 mutant.

**Figure 2-figure supplement 3.**
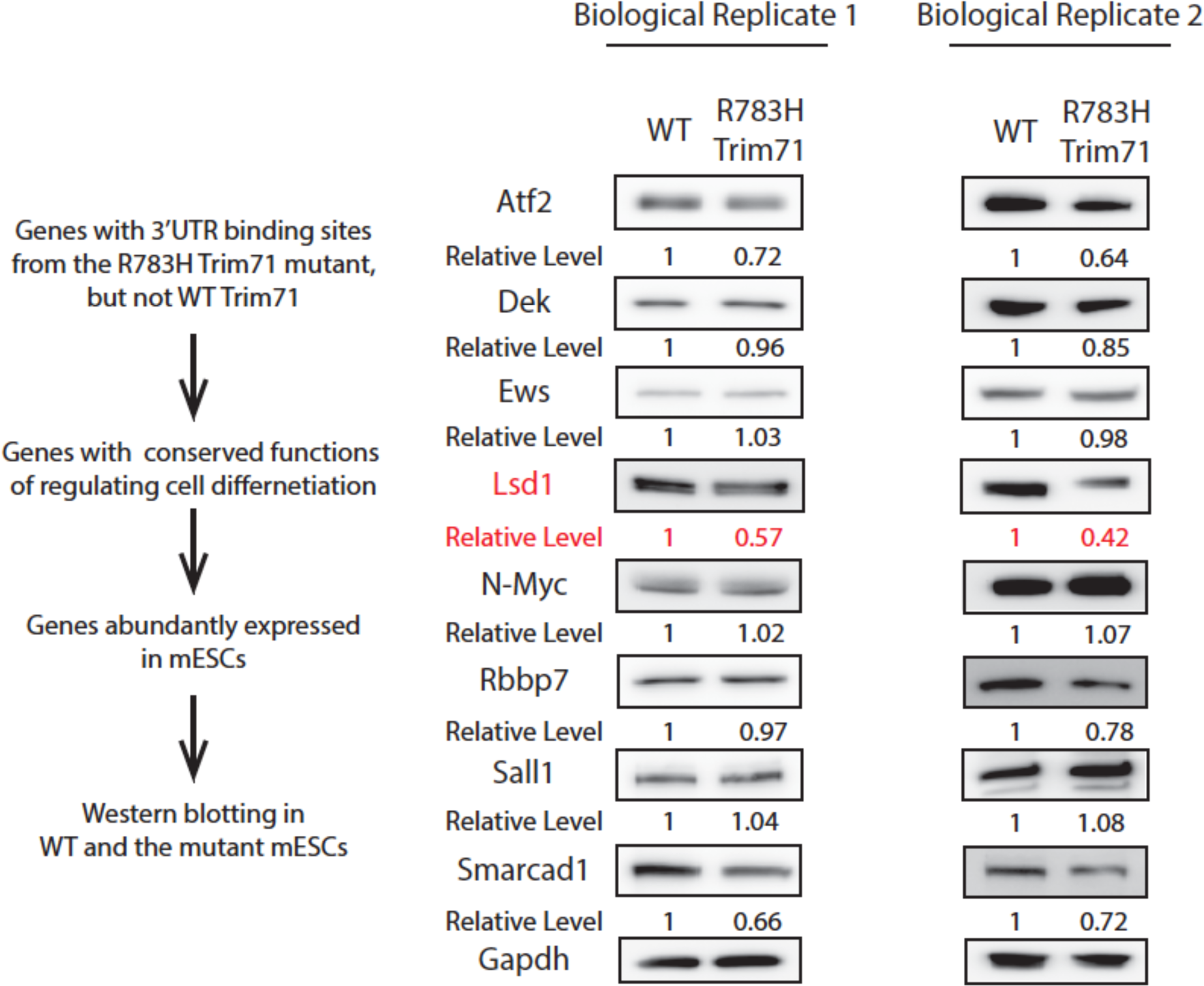
Identification of potential functional targets of the R783H Trim71 mutant. In each biological replicate of Western blotting, Gapdh was used for normalization in calculating the relative levels.

**Figure 3-figure supplement 1.**
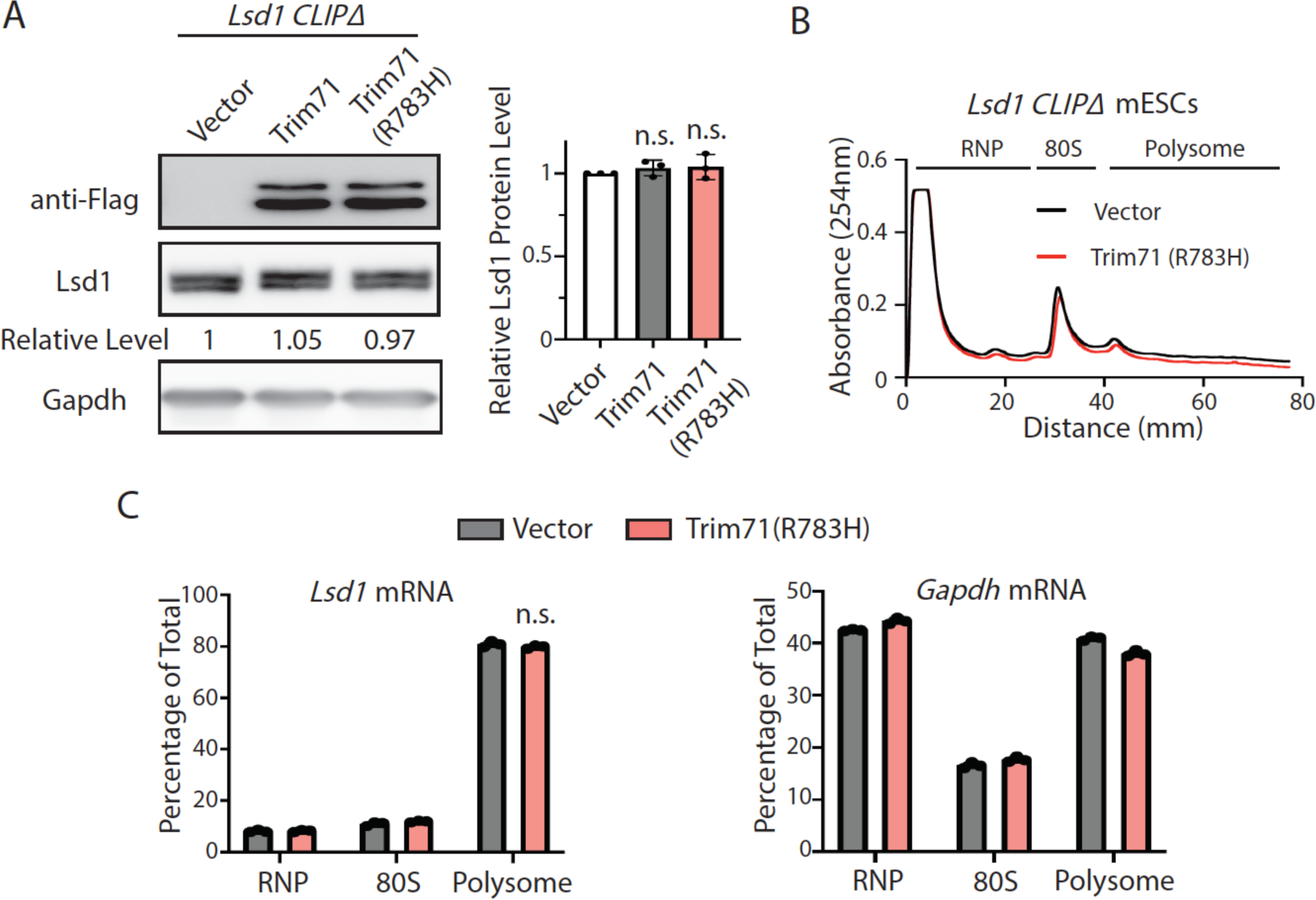
Repression of *Lsd1* mRNA translation by the Trim71 mutant R783H is dependent on its binding to *Lsd1* mRNA. A. A representative Western blot of the CLIPΔ mESCs expressing an empty vector, Flag-Trim71, and Flag-Trim71(R783H). Gapdh was used as a loading control for quantification of Lsd1 levels. The quantification results represent the means (± SD) of three independent experiments. n.s. not significant (p>0.05) by the Student’s t-test. B. Polysome analysis in the CLIPΔ mESCs expressing an empty vector and Flag- Trim71(R783H). C. Quantification of the indicated mRNA distribution in the RNP, 80S, and polysome fractions from the CLIPΔ mESCs expressing an empty vector and Flag- Trim71(R783H). The results represent the means (± SD) of three independent experiments. n.s. not significant (p>0.05) by the Student’s t-test.

**Figure 3-figure supplement 2.**
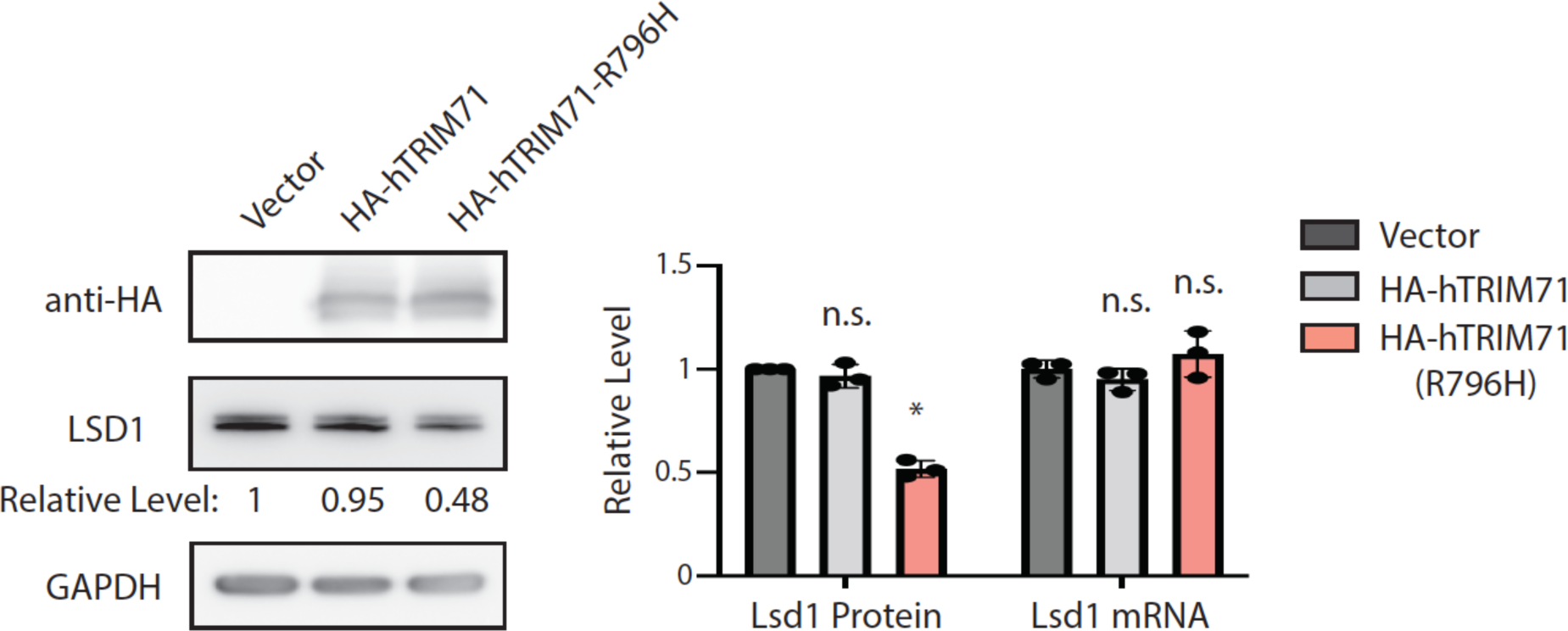
The human TRIM71 mutant R796H represses LSD1. A representative Western blotting the NCCIT cells expressing an empty vector, HA- hTRIM71, and HA-hTrim71(R796H). Gapdh was used for normalization in the quantification of Lsd1 protein levels, and 18S rRNA was used for normalization in the qRT-PCR quantification of Lsd1 mRNA levels. The quantification results represent the means (± SD) of three independent experiments. * p < 0.05, n.s. not significant (p>0.05) by the Student’s t-test.

**Figure 4-figure supplement 1.**
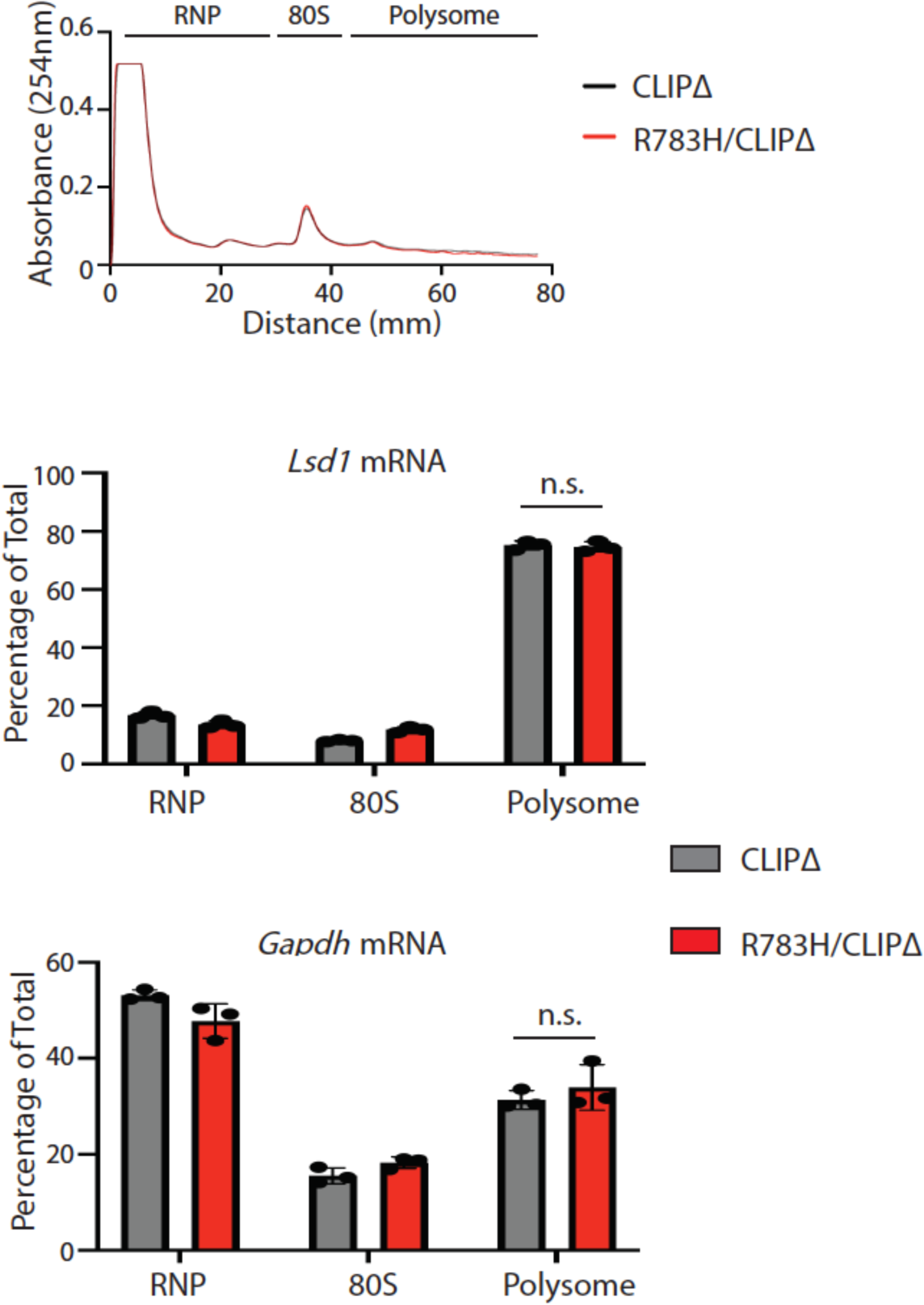
The R783H mutation in the Lsd1 CLIPΔ background does not alter the polysome association of Lsd1 mRNA. Polysome analysis was performed in the Lsd1 CLIPΔ mESCs and the Lsd1 CLIPΔ/R783H mESCs. qRT-PCR was used to quantified the indicated mRNAs in the RNP, 80S, and polysome regions on the sucrose density gradient. The results represent the means (± SD) of three independent experiments. n.s. not significant (p>0.05) by the Student’s t-test.

**Figure 5-figure supplement 1.**
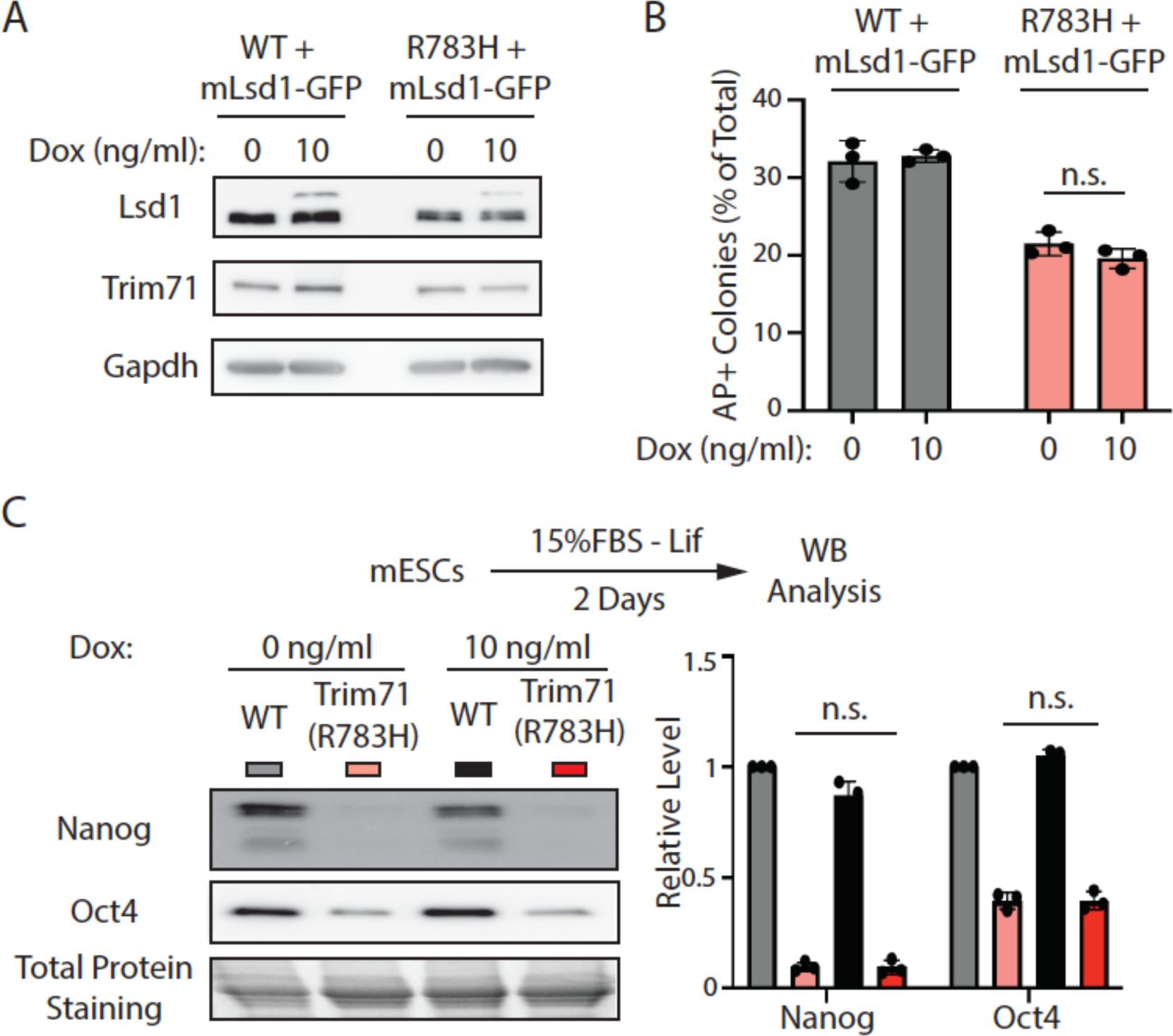
The demethylase catalytic mutant Lsd1 fails to alleviate the stem cell differentiation defects in the Trim71(R783H) mESCs. A. Western blotting in the WT and the Trim71(R783H) mESCs with dox-inducible expression of a demethylase catalytic mutant (K661A) Lsd1 (mLsd1-GFP). B. Exit pluripotency assay for mESCs. C. Representative Western blotting and quantification of pluripotency factors during the monolayer differentiation of mESCs. In B, the colony morphology and AP intensity were evaluated through microscopy. 100- 200 colonies were examined each time to determine the percentage of undifferentiated colonies. The quantification results from B and C represent the means (± SD) of three independent experiments. Total protein levels were used for normalization in the quantification results of C. *p<0.05; and n.s. not significant (p>0.05) by the Student’s t- test.

